# BioIMA: a one-click desktop tool for standardized extraction of phenotypic traits from biological images

**DOI:** 10.64898/2026.08.30.747465

**Authors:** Xiner Qumu, Xuming Dan, Jianju Feng, Yangkai Cui, Yi Gong, Yukang Hou, Qiang Lai, Zeng Wang, Yulin Zhang, Yiman Zhu, Yan Yu, Feng Zhang, Marco Todesco, Jing Wang

## Abstract

Standardized extraction of quantitative phenotypes from images is increasingly important across plant biology, from ecological and evolutionary studies to genetics, breeding, and functional genomics. However, as large image datasets are increasingly used for trait analysis, many biologically relevant traits, including size, shape, color, and spatial patterning, are still measured manually or using fragmented semi-automated workflows. These limitations reduce throughput, reproducibility, and accessibility, especially for researchers without computational expertise. Here, we present BioIMA, an open-source desktop tool for rapid and standardized phenotyping from biological images. BioIMA integrates foundation model-based segmentation with automated trait computation, allowing users to extract quantitative measurements from images through an intuitive graphical interface and without model training. To validate its performance, we quantified a set of knot morphological traits in two *Populus* species, as these measurements are typically time-consuming to perform manually. Automatic measurements showed strong agreement with manual ImageJ-based measurements (*R*² > 0.95), while reducing per-image processing time by approximately 75% (from ∼15 s to ∼4 s). BioIMA was further applied to diverse plant datasets, including *Helianthus* and *Rhododendron* images with varying morphologies and background conditions. Although developed for plant phenotyping, BioIMA may also be extended to other biological samples where region-based size, shape, or color traits are of interest. By combining accessibility and standardization in a lightweight local application, BioIMA provides a practical community resource for image-based phenotyping in ecological and evolutionary studies.

## Introduction

Image-based phenotyping has become an essential approach in ecological, and evolutionary research, enabling quantitative characterization of traits such as size, shape, color, and pattern across increasing spatial and temporal scales (Jiang & Li, 2020; Murphy et al., 2024). The rapid development of imaging technologies has greatly expanded the capacity to collect phenotypic data. However, extracting biologically meaningful information from these images remains a major bottleneck (Furbank & Tester, 2011; Gill et al., 2022). This constraint persists as phenotypic measurements often lag behind advances in genomics and data acquisition, limiting our ability to link genotype, phenotype, and environment. With the increasing demand for various types of phenotypic measurements in plant sciences, there is a growing need for customized tools and approaches to rapidly extract biologically relevant information from images.

Despite progress in imaging technologies, trait extraction in plant, ecological, and evolutionary research still relies heavily on manual measurement or fragmented semi-automated workflows, e.g., combining ImageJ-based segmentation with separate scripts for measurement and statistical export (Gehan et al., 2017; Schneider et al., 2012). Manual phenotyping is labor-intensive, time-consuming, and prone to observer bias, particularly when applied to complex morphologies or moderate to large sample sizes (Fahlgren et al., 2015). While dedicated image analysis tools exist, many require programming expertise that limits their adoption among biologists without computational backgrounds (Pieruschka & Schurr, 2019). As a result, the transition from raw images to analyzable trait data remains a practical bottleneck even in research groups that already have imaging infrastructure in place.

Recent advances in foundation models for computer vision, such as the Segment Anything Model (SAM), provide new opportunities for flexible and generalizable image segmentation (Kirillov et al., 2023). These models enable prompt-based and zero-shot (i.e., not requiring dedicated training) segmentation across diverse image domains, reducing the need for task-specific training and annotated datasets. However, existing applications of such models in plant and ecological research have largely focused on segmentation or annotation tasks alone (Abbey & Meroz, 2026; Williams et al., 2024). As a result, a gap remains between advances in segmentation models and practical tools for extracting biologically meaningful, analysis-ready phenotypic data, such that even researchers who adopt SAM-based segmentation must still rely on separate, often manual, steps to extract and organize trait data.

Here we present BioIMA, an open-source, one-click desktop application designed to bridge the gap between image segmentation and quantitative trait extraction. To demonstrate its utility, we used tree knot morphology as a representative use case. Knot morphology is closely related to wood quality, but its quantitative measurement is often labor-intensive and time consuming when performed manually. This has motivated continued efforts toward automated measurement and detection in forestry research (Duchateau et al., 2013; Roussel et al., 2014). Furthermore, we evaluated BioIMA on diverse plant image datasets encompassing varied morphologies and imaging conditions, supporting its broad applicability to any dataset where region-based traits such as size, shape, or color are of interest and require measurement.

## Materials and methods

### Tool Architecture and Implementation

BioIMA is implemented using the Avalonia UI framework, which enables cross platform desktop deployment on Windows and macOS through a unified graphical interface (Figure 1). The software is distributed as a self-contained installable package with all required dependencies, allowing local use without manual configuration of deep learning environments. Image segmentation is powered by embedded models currently including SAM (Kirillov et al., 2023) and mobile SAM (Zhang et al., 2023), which are executed locally through ONNX Runtime for efficient inference without internet connectivity. Source code, documentation, example datasets, and a user manual are publicly available on GitHub (https://github.com/jingwanglab/BioIMA).

**Figure 1.**
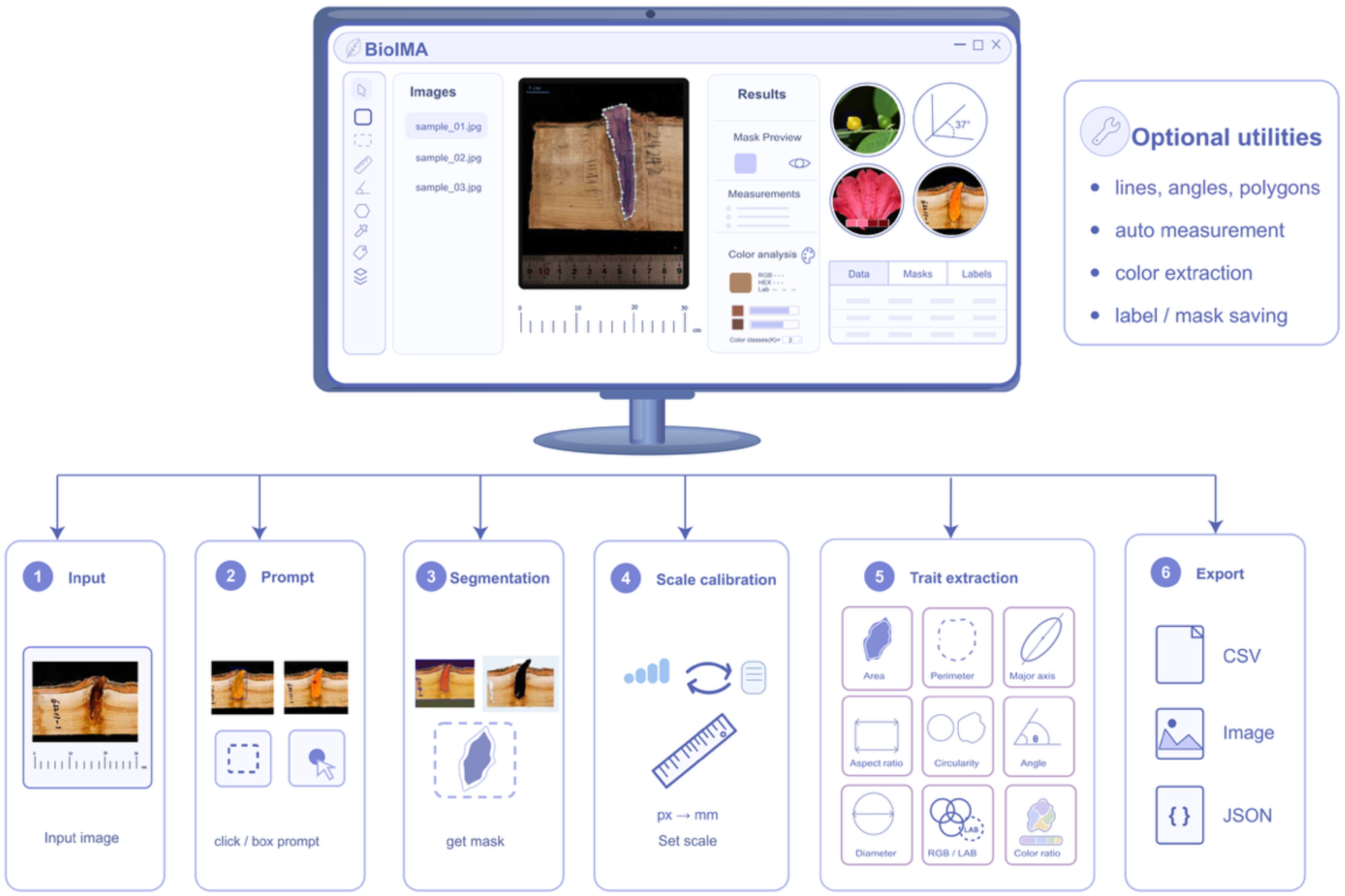
Overview of the BioIMA workflow and interaction options. BioIMA supports prompt-based SAM segmentation with optional manual refinement to obtain a finalized mask for each region of interest. The finalized mask is scale-calibrated and used to derive quantitative morphological traits, including area, perimeter, shape descriptors, RGB and CIE Lab values, HEX color codes and color distribution summaries. Results are exported as a CSV table and color summary images, and masks/labels can be saved for downstream use.

### Input and automatic segmentation processing pipeline

BioIMA accepts standard image formats, including JPEG, PNG, and TIFF, and is designed to process biological images with complex backgrounds or subtle object boundaries. After image loading, users initiate segmentation with a lightweight prompt, either a click indicating the target region or an interactively defined bounding box. The embedded model then generates a pixel-level mask for the region of interest, which can be reviewed immediately. When necessary, the mask can be refined by adding additional prompts or through manual polygon editing. This interactive workflow allows segmentation results to be adjusted before downstream trait extraction.

### Scale calibration and trait extraction

All morphological measurements are computed from the finalized binary mask. To report traits in real units, BioIMA supports scale calibration tailored to each image using an in-frame scale bar. Users define a reference line on a known length (e.g., a scale bar) and provide its true value, allowing conversion from pixel units to physical units (Schneider et al., 2012).

Then, the tool performs instant phenotyping processing to extract relevant morphological traits, including area, perimeter, major axis length, aspect ratio, circularity metrics, equivalent diameter, and line-based measurements (length and angle) (Bradski, 2000; Gehan et al., 2017). In addition, color features are computed in both RGB and CIE Lab spaces, including mean RGB values, Lab coordinates, and HEX color codes. BioIMA also includes an optional color correction step that uses a user selected neutral white or gray reference region to partially reduce color bias caused by lighting, camera settings, or scanning conditions. Subsequent color measurements and color distribution analyses are performed on the corrected image. For regions with heterogeneous or patterned coloration, BioIMA provides a color distribution module that uses k means clustering to partition pixels within the selected region into dominant color classes. Users can specify the number of color clusters according to their analysis needs, and BioIMA then estimates the relative proportion of each color class. The resulting color summary can be exported as a report, allowing users to inspect representative colors, color proportions, and region level color statistics. These traits are calculated using standard computer vision operations (mask binarization, contour extraction, and shape descriptors based on boundaries) to ensure interpretability and reproducibility (Bradski & Kaehler, 2000; Van der Walt et al., 2014). All outputs are compiled into structured phenotype tables (CSV format), enabling direct integration with downstream statistical, ecological, or genomic analyses. Segmentation masks can also be exported for reinspection or reuse.

### Dataset and validation

We evaluated BioIMA on a knot image dataset from two desert poplar species, *Populus euphratica* and *Populus alba* var. *pyramidalis*, which were collected in March 2023 in Xinjiang, China. These species represent ecologically important woody plants in arid regions (Guo et al., 2025; Ma et al., 2018), and knot morphology in their stems provides a tractable case for validating image-based trait extraction given its discrete boundaries and measurable geometry. Images were acquired using a SONY A7RM3 camera equipped with a Sigma 24-70 mm F2.8 DG DN ART lens under standardized lighting and background conditions, with a physical scale bar included in each frame (see image acquisition section; Figure S1). Measurement validity was assessed by comparison with manual measurements obtained in ImageJ (Schneider et al., 2012) as a reference standard. For each image, knot area, major axis length, and perimeter were measured independently: once using BioIMA’s automated segmentation pipeline, and once by a single trained observer performing manual polygon annotation in ImageJ. Agreement between methods was evaluated using linear regression (*R*²) and Bland-Altman analysis (Bland & Altman, 1986) to quantify systematic bias and limits of agreement. Operational efficiency and repeatability were assessed on a randomly selected subset of images (n = 100). Each image was measured five times by the same observer with each method. All statistical analyses were performed in R (v4.1.2) (R Core Team, 2024).

## Results

### Interface Overview and Workflow

BioIMA combines image visualization, mask inspection, scale calibration, trait measurement, and data export within a single graphical workspace (Figure 2). For each selected region, the interface displays an overlaid mask preview and calibrated measurements, allowing users to inspect segmentation quality before exporting trait data. The tool supports two workflows for trait extraction. In the automatic mode, SAM-based interactive segmentation generates a region mask from minimal user input, such as a single click or bounding box. Users can review and refine the mask and accept it, after which morphological traits are computed immediately from the finalized mask (Figure 2a, b). In the manual mode, users define regions using drawing tools such as lines and polygons, and the same measurement pipeline is applied to the annotated region. When automated boundaries require correction, or when manual delineation is preferred, users can refine masks or define regions. Scale calibration, color summarization, and CSV export are integrated into the same workflow to support standardized trait extraction and downstream reuse (Figure 2c-e), The exported trait fields and their definitions are provided in Table S1.

**Figure 2.**
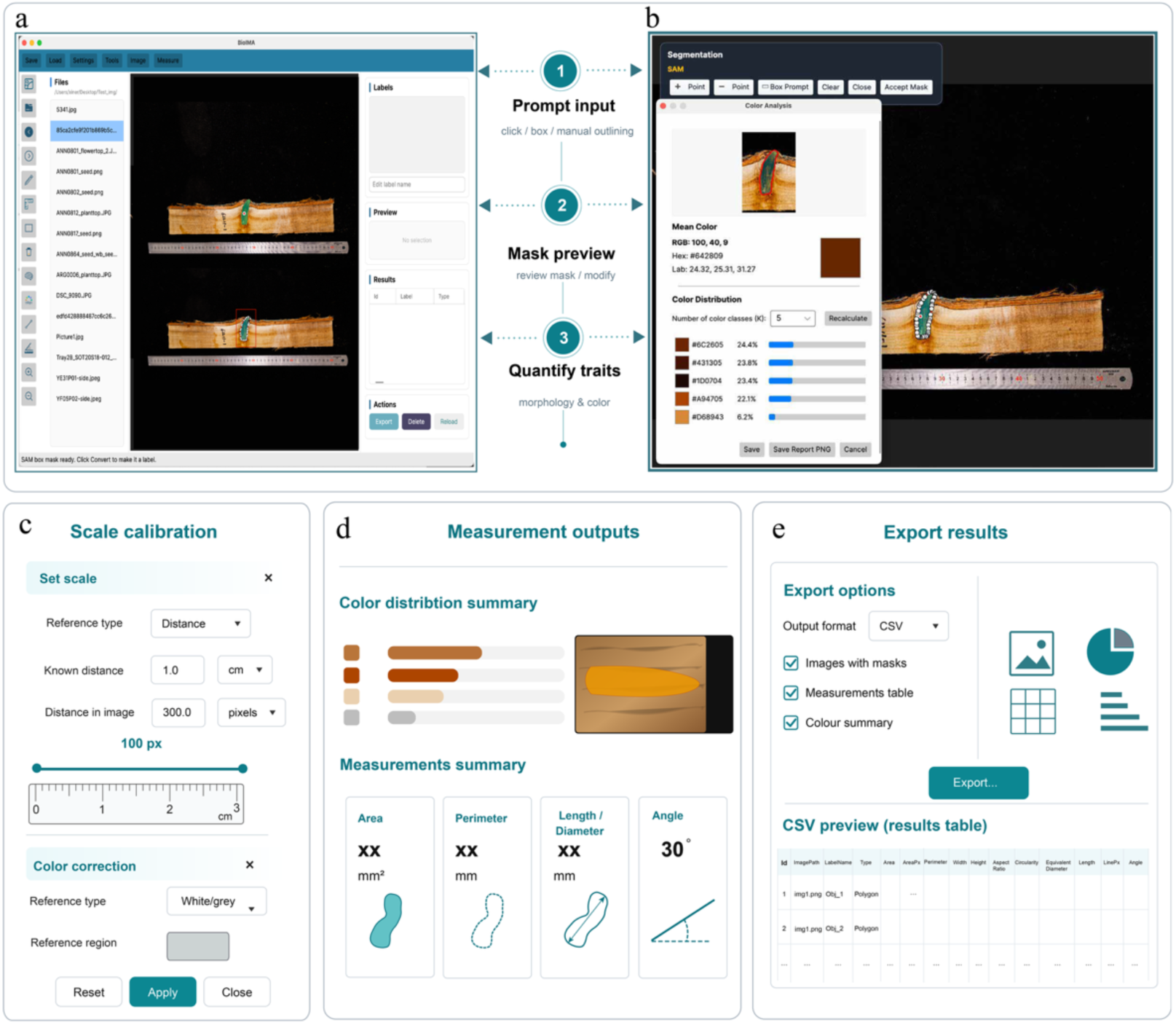
BioIMA workflow and representative analysis outputs. (a) Users generate segmentation masks from point, box, or manual outline prompts and review the resulting masks interactively. (b) After mask confirmation, BioIMA computes quantitative traits from the selected region, including morphological measurements and color analysis summaries. The color analysis module reports RGB, HEX, and CIE Lab values together with color distribution profiles that can be adjusted using different cluster numbers (k). (c) Scale calibration converts pixel measurements into physical units using an in-frame reference, while color correction reduces color bias using a white or gray reference region. (d) Representative quantitative outputs, including morphological measurements and color distribution summaries derived from finalized masks. (e) Export module showing structured CSV outputs, color summary images, and measurement records for downstream analysis.

### Accuracy, agreement, and throughput benchmarking

Measurement agreement between BioIMA and ImageJ was evaluated using the knot image dataset from two *Populus* species. Representative examples illustrate the corresponding BioIMA segmentation and ImageJ manual delineation of the same knot region (Figure 3a, b). We compared three traits derived from masks (knot area, perimeter, and major axis length) between BioIMA and ImageJ measurements (Figure 3c-e). Across all three traits, BioIMA measurements showed strong linear agreement with ImageJ, with coefficients of determination of *R*²= 0.99 for area and *R*²= 0.96 for both perimeter and length. Bland-Altman analysis of knot area measurements further indicated limited systematic bias, with most observations falling within the 95% limits of agreement (Figure 3f). Visual inspection suggested that the few outlying cases were mainly associated with larger knots or visually ambiguous boundaries.

**Figure 3.**
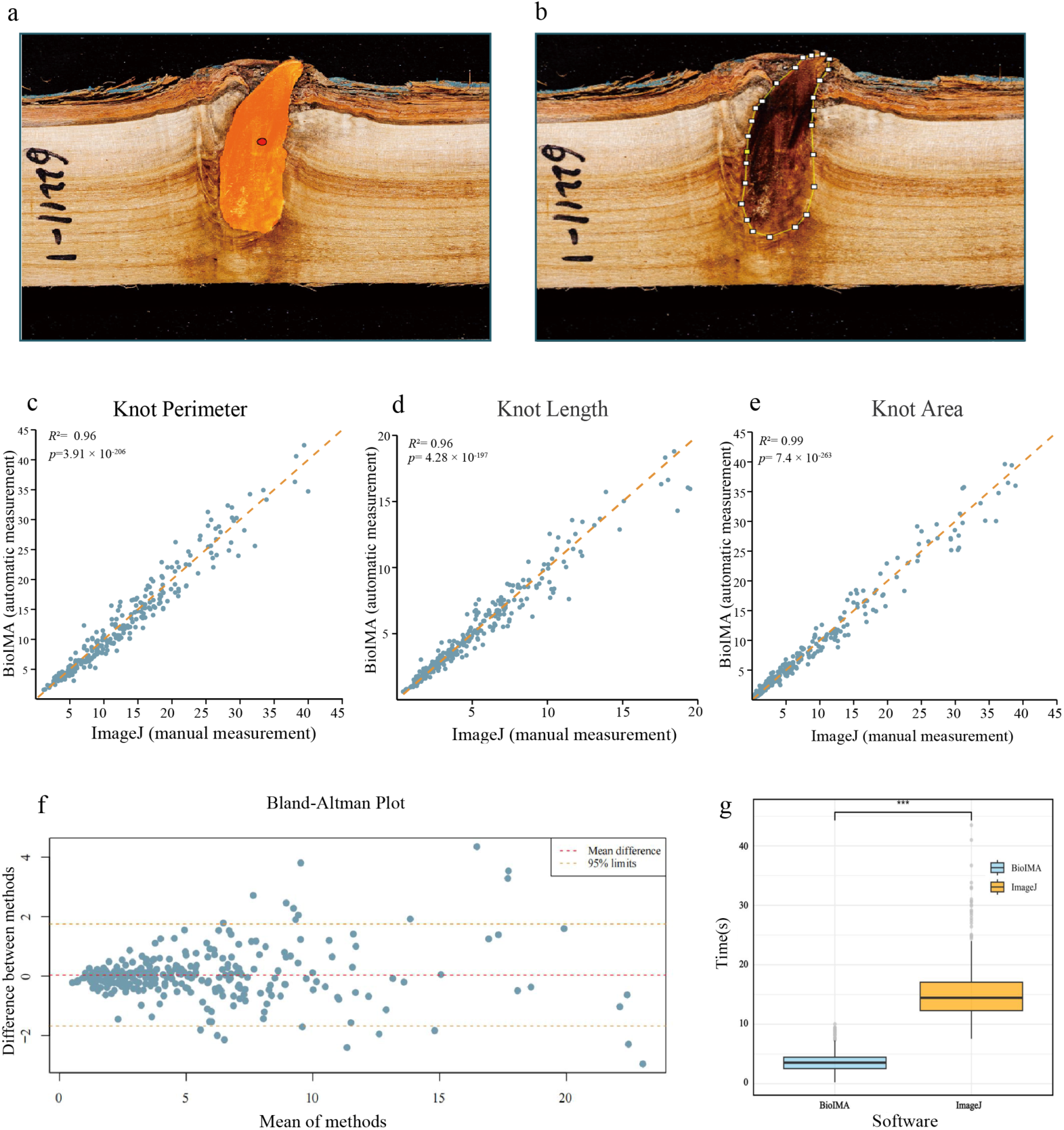
Benchmarking BioIMA with ImageJ on knot measurements. (a) Representative knot region segmented using BioIMA. (b) Corresponding knot region manually delineated in ImageJ. (c-e) Agreement between BioIMA and ImageJ measurements for knot perimeter (c), major axis length (d), and area (e). The dashed line indicates the 1:1 reference line, and *R*² values are shown for each trait. (f) Bland-Altman analysis of knot area measurements, showing the mean difference and 95% limits of agreement. (g) Comparison of per-image measurement time between BioIMA and ImageJ. Boxplots show the distribution of measurement times across repeated measurements, with BioIMA reducing median measurement time by 75.5%.

We next quantified throughput using a repeated measurement experiment on 100 knot images measured five times each (*N* = 500 per method). ImageJ exhibited a broader time distribution (median 14.445 s; IQR 12.284-17.069 s), whereas BioIMA achieved substantially faster and more consistent processing times (median 3.534 s; IQR 2.530-4.454 s) (Figure 3g), corresponding to a 75.5% reduction in median time per image. Raw measurements and replicate timing data for this benchmark are provided in supplementary Table S2.

### Application to diverse plant organs and imaging conditions

To illustrate BioIMA’s applicability beyond the knot dataset used for benchmarking, we applied the same prompt-guided workflow to representative plant images spanning different organs and acquisition conditions (Figure 4). These examples included woody structures, leaves, seeds, flowers, fruits, belowground tissues, and a non-plant biological structure with measurements including region segmentation, color summary, size and shape traits, and line or angle based traits. For images containing multiple objects, regions were selected and measured sequentially rather than through fully automated multi object detection. Using the same comparison framework as in the knot benchmark, we further evaluated BioIMA on *Helianthus* inflorescence and *Rhododendron* petal datasets (Figure 4a, b). BioIMA showed high agreement with ImageJ for area measurements in both datasets (*R*² = 0.93-0.97; Fig. S2), supporting application of the same workflow beyond knot images. Raw measurements used for this additional validation are provided in supplementary Table S3. Together, these examples show that BioIMA can be applied across a broad range of plant organs and imaging contexts while maintaining a consistent workflow for trait extraction and export.

**Figure 4.**
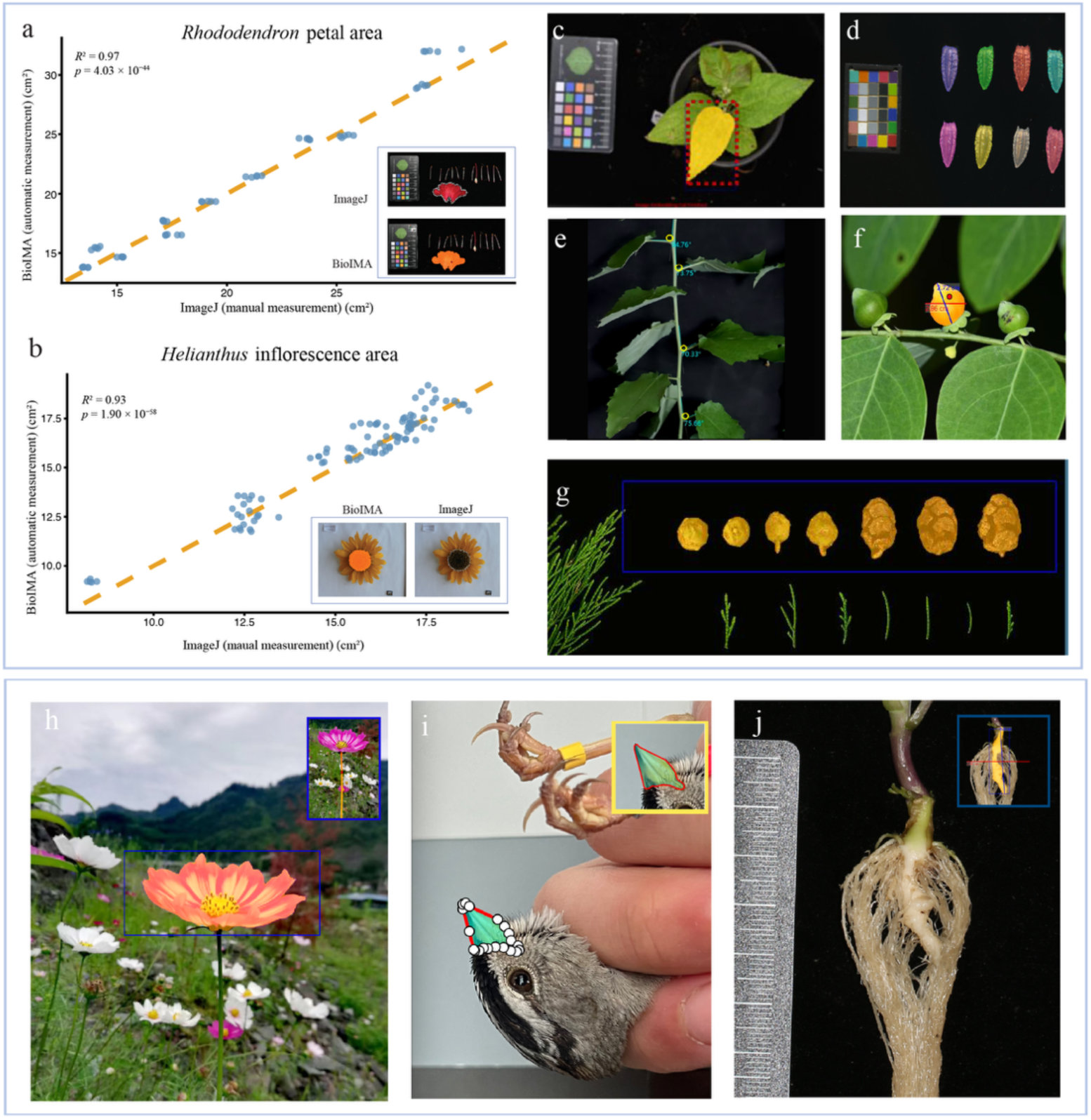
Examples of BioIMA applied to diverse plant organs and other biological samples. (a,b) Additional comparisons between BioIMA and ImageJ for area measurements in *Rhododendron* petals and *Helianthus* inflorescences, each image was measured five times; the *Rhododendron* validation included 10 images (50 measurements total), and the *Helianthus* validation included 20 images (100 measurements total). Scatter plots show agreement between methods, with dashed lines indicating the identity relationship; representative original and segmented examples are shown as insets. (c-j) Representative applications of the same workflow to user selected regions across different imaging contexts and trait types. Examples include (c) leaf region segmentation using a box prompt, (d) sequential measurement of multiple seeds within a single image, (e) line and angle measurements on leaves, (f) fruit size measurement, (g) sequential size measurement of cone like reproductive structures, (h) flower region and stem length segmentation under a complex field background, (i) color distribution analysis of a patterned bird beak region as an example of applying BioIMA to non-plant biological samples. (j) Measurement of clubroot associated root swelling in *Orychophragmus violaceus.* The selected region illustrates how BioIMA can support delineation and quantification of disease-related root structures with low color contrast from surrounding tissue.

## Discussion

Plant genomics has entered a big data era, yet a persistent challenge in plant and ecological research is that the generation of genomic and environmental data often outpaces the acquisition of standardized phenotypic measurements (Furbank & Tester, 2011; Tardieu et al., 2017). This gap is particularly acute when trait acquisition relies on manual measurement, which remains labor-intensive, sensitive to observer bias, and difficult to standardize across large or heterogeneous datasets (Giuffrida et al., 2018). Image-based phenotyping provides an important route for scaling trait acquisition, but many existing workflows still require substantial manual annotation, parameter adjustment, or programming before image regions can be converted into quantitative measurements suitable for downstream analyses.

BioIMA was developed to address this practical gap. It uses prompt-guided segmentation as an intermediate step and converts selected image regions directly into calibrated morphological and color traits. This design is useful for biological images in which boundaries are difficult to define automatically, such as woody tissues, overlapping organs, field photographs, or samples with heterogeneous backgrounds (Okyere et al., 2023; Zenkl et al., 2022). By integrating locally executed segmentation, scale calibration, interactive mask inspection, and direct export of quantitative trait tables and color distribution reports within a graphical interface, BioIMA streamlines image-based phenotyping and reduces the time and expertise required to generate standardized image derived traits.

The benchmarking results suggest that BioIMA can produce measurements that are highly consistent with ImageJ-based manual measurement while substantially improving efficiency. Across multiple traits measures on different tissue types (wood knots, petals, inflorescences), BioIMA showed strong agreement with ImageJ, indicating that prompt-guided segmentation can provide reliable quantitative measurements for this use case. The Bland-Altman analysis further suggested limited systematic bias for most samples, although larger or visually ambiguous features (e.g., wood knots) remained more prone to deviation. This pattern is expected because ambiguous boundaries affect both manual and automated measurements, and it highlights the value of keeping users in the loop for mask inspection and correction (Pachitariu & Stringer, 2022). The reduction in median processing time by 75.5% further supports the potential of BioIMA for studies that require repeated or large-scale trait extraction.

Although the main validation focused on knot morphology in two *Populus* species, the broader prompt and measure workflow is not restricted to this trait. The additional examples across leaves, flowers, fruits, seeds, roots, and other plant structures illustrate that BioIMA can be applied to diverse regions of interest when an interpretable mask can be obtained. The color distribution example on a patterned bird beak further suggests that the same region-based workflow may be extended to non-plant biological structures where localized color or pattern traits are of interest. These examples should be viewed as demonstrations of applicability rather than full quantitative validation for every organ or imaging condition. Additional benchmarking on specific traits, species, and imaging environments will be important for future studies, especially when BioIMA is used for highly complex backgrounds, overlapping objects, or traits with subtle color and texture variation (Jiang & Li, 2020). Nevertheless, while the current implementation supports local deployment and avoids the need for model training, performance still depends on image quality, object contrast, and the suitability of the selected prompt. Therefore, clear image acquisition protocols and routine mask inspection remain essential for reproducible use. To improve scalability while addressing the current reliance on user-guided target selection, we are incorporating automated instance detection and batch processing functions for large image collections. These functions are intended to reduce manual effort while preserving user oversight, which remains important because fully automatic segmentation can be unreliable in complex biological scenes (Kirillov et al., 2023; Okyere et al., 2023).

Overall, BioIMA provides a practical resource for converting biological images into standardized phenotype tables through an accessible desktop workflow. By integrating foundation model-based segmentation, scale-calibrated measurement, interactive quality control, and structured data export within a single interface, BioIMA streamlines quantitative phenotyping and reduces the time and expertise required to generate analysis-ready trait data. As biological image collections continue to expand, tools that make trait extraction more accessible and reproducible will be increasingly important for linking phenotypes with genomic, environmental, and experimental data across plant, ecological, and evolutionary research.

## Supporting information

Supplementary Table S2: Raw Populus knot measurement and timing benchmark data

Supplementary Table S3: Raw Helianthus and Rhododendron validation data

BioIMA User Manual (English)

## Author contributions

J.W. and X.Q. conceived the study. X.Q., X.D., J.F., Y.C., Y.G., Y.H., Q.L. Z.W., Y.Z., Y.Z. handled the sampling and analyzed the knot data. X.Q. developed the BioIMA tool. X.Q., X.D., and J.W wrote the manuscript, with revision from M.T. All authors approved the final version of the manuscript.

## Data availability

All data required to evaluate the conclusion from the study are provided within the main text and/or the supplementary materials. The BioIMA and the associated manual are provided at https://github.com/jingwanglab/BioIMA.

## Acknowledgements

This work was supported by the National Key Research and Development Program of China (2022YFD2201200) to J.W.

## Figures and tables

**Figure S1 | Preparation and representative examples of the *Populus* knot image dataset used for validation.**

(a,b) Field sampling and collection of felled log materials. (c,d) Preparation of knot containing plank samples. (e,f) Representative validation images from *Populus euphratica* (e) and *Populus alba* var. *pyramidalis* (f), which were used for comparison between BioIMA and ImageJ measurements. Images include in frame scale references for calibration.

**Figure S2 | Additional validation of BioIMA area measurements on *Helianthus* inflorescence and *Rhododendron* petal images.**

BioIMA area measurements were compared with ImageJ manual measurements using independent *Rhododendron* petal and *Helianthus* inflorescence image datasets. The left panels show BioIMA-ImageJ measurement agreement, with orange dashed lines indicating the 1:1 identity line; *R*² and *p* values are shown in each panel. The right panels show Bland-Altman analyses, with red dashed lines indicating mean bias and yellow dashed lines indicating the 95% limits of agreement. Both datasets showed high agreement between methods

**Table S1 | BioIMA exported trait fields and definitions.** List of trait fields exported by BioIMA in CSV format, including label metadata, morphometric measurements, RGB/HEX/Lab color summaries, and color distribution outputs.

**Table S2 | Raw *Populus* knot measurement and timing benchmark data.** Raw BioIMA and ImageJ benchmark data for *Populus* knot images. The first sheet contains measurement time records used for the software throughput comparison in Fig. 3g. The second sheet contains paired morphometric measurements used for the method-comparison and Bland-Altman analyses in Fig. 3c-f, including knot perimeter, major axis length, and knot area. (excel)

**Table S3 | Raw *Helianthus* and *Rhododendron* validation data.** Raw BioIMA and ImageJ measurements for *Helianthus* inflorescence and *Rhododendron* petal area validation, including sample identifiers and measurement values used to generate Fig. S2. (excel)

## References

Abbey, A., & Meroz, Y. (2026). Segment any plant (SAP): Foundation-model segmentation for plant time-series phenotyping [Preprint]. bioRxiv. 10.64898/2026.03.11.711099

Bland, J. M., & Altman, D. (1986). Statistical methods for assessing agreement between two methods of clinical measurement. The Lancet, 327(8476), 307–310.

Bradski, G., & Kaehler, A. (2000). OpenCV. Dr. Dobb’s Journal of Software Tools, 3(2), 1–81.

Duchateau, E., Longuetaud, F., Mothe, F., Ung, C., Auty, D., & Achim, A. (2013). Modelling knot morphology as a function of external tree and branch attributes. Canadian Journal of Forest Research, 43(3), 266–277. 10.1139/cjfr-2012-0365

Fahlgren, N., Gehan, M. A., & Baxter, I. (2015). Lights, camera, action: High-throughput plant phenotyping is ready for a close-up. Current Opinion in Plant Biology, 24, 93–99.

Furbank, R. T., & Tester, M. (2011). Phenomics technologies to relieve the phenotyping bottleneck. Trends in Plant Science, 16(12), 635–644.

Gehan, M. A., Fahlgren, N., Abbasi, A., Berry, J. C., Callen, S. T., Chavez, L., Doust, A. N., Feldman, M. J., Gilbert, K. B., & Hodge, J. G. (2017). PlantCV v2: Image analysis software for high-throughput plant phenotyping. PeerJ, 5, e4088.

Gill, T., Gill, S. K., Saini, D. K., Chopra, Y., De Koff, J. P., & Sandhu, K. S. (2022). A comprehensive review of high-throughput phenotyping and machine learning for plant stress phenotyping. Phenomics, 2(3), 156–183. 10.1007/s43657-022-00048-z

Giuffrida, M. V., Chen, F., Scharr, H., & Tsaftaris, S. A. (2018). Citizen crowds and experts: Observer variability in image-based plant phenotyping. Plant Methods, 14(1), 12. 10.1186/s13007-018-0278-7

Guo, X., Li, J., Zhang, J., Wei, C., & Li, Z. (2025). Vegetation growth improvement Inadequately represents the ecological restoration of the *Populus euphratica* forests in Xinjiang, China. Ecological Indicators, 170, 113086. 10.1016/j.ecolind.2025.113086

Jiang, Y., & Li, C. (2020). Convolutional neural networks for image-based high-throughput plant phenotyping: A review. Plant Phenomics, 2020, 4152816. 10.34133/2020/4152816

Kirillov, A., Mintun, E., Ravi, N., Mao, H., Rolland, C., Gustafson, L., Xiao, T., Whitehead, S., Berg, A. C., Lo, W.-Y., Dollar, P., & Girshick, R. (2023). Segment anything. In Proceedings of the IEEE/CVF International Conference on Computer Vision (pp. 4015–4026). https://openaccess.thecvf.com/content/ICCV2023/html/Kirillov_Segment_Anything_ICCV_2023_paper.html

Ma, J., Wan, D., Duan, B., Bai, X., Bai, Q., Chen, N., & Ma, T. (2018). Genome sequence and genetic transformation of a widely distributed and cultivated poplar. Plant Biotechnology Journal, 17(2), 451–460. 10.1111/pbi.12989

Murphy, K. M., Ludwig, E., Gutierrez, J., & Gehan, M. A. (2024). Deep Learning in Image-Based Plant Phenotyping. Annual Review of Plant Biology, 75(1), 771–795. 10.1146/annurev-arplant-070523-042828

Okyere, F. G., Cudjoe, D., Sadeghi-Tehran, P., Virlet, N., Riche, A. B., Castle, M., Greche, L., Mohareb, F., Simms, D., & Mhada, M. (2023). Machine learning methods for automatic segmentation of images of field-and glasshouse-based plants for high-throughput phenotyping. Plants, 12(10), 2035.

Pachitariu, M., & Stringer, C. (2022). Cellpose 2.0: How to train your own model. Nature Methods, 19(12), 1634–1641.

Pieruschka, R., & Schurr, U. (2019). Plant Phenotyping: Past, Present, and Future. Plant Phenomics, 2019, 7507131. 10.34133/2019/7507131

R Core Team. (2024). R: A Language and Environment for Statistical Computing. R Foundation for Statistical Computing. https://www.R-project.org/

Roussel, J.-R., Mothe, F., Krähenbühl, A., Kerautret, B., Debled-Rennesson, I., & Longuetaud, F. (2014). Automatic knot segmentation in CT images of wet softwood logs using a tangential approach. Computers and Electronics in Agriculture, 104, 46–56.

Schneider, C. A., Rasband, W. S., & Eliceiri, K. W. (2012). NIH Image to ImageJ: 25 years of image analysis. Nature Methods, 9(7), 671–675.

Tardieu, F., Cabrera-Bosquet, L., Pridmore, T., & Bennett, M. (2017). Plant phenomics, from sensors to knowledge. Current Biology, 27(15), R770–R783.

Van der Walt, S., Schönberger, J. L., Nunez-Iglesias, J., Boulogne, F., Warner, J. D., Yager, N., Gouillart, E., & Yu, T. (2014). scikit-image: Image processing in Python. PeerJ, 2, e453.

Williams, D., Macfarlane, F., & Britten, A. (2024). Leaf only SAM: A segment anything pipeline for zero-shot automated leaf segmentation. Smart Agricultural Technology, 8, 100515.

Zenkl, R., Timofte, R., Kirchgessner, N., Roth, L., Hund, A., Van Gool, L., Walter, A., & Aasen, H. (2022). Outdoor plant segmentation with deep learning for high-throughput field phenotyping on a diverse wheat dataset. Frontiers in Plant Science, 12, 774068.

Zhang, C., Han, D., Qiao, Y., Kim, J. U., Bae, S.-H., Lee, S., & Hong, C. S. (2023). Faster segment anything: Towards lightweight SAM for mobile applications [Preprint]. arXiv. 10.48550/arXiv.2306.14289

