## Supplementary Figures and Tables for "BioIMA: a one-click desktop tool for standardized extraction of phenotypic traits from biological images"

**Figure S1 | Preparation and representative examples of the *Populus* knot image dataset used for validation.**
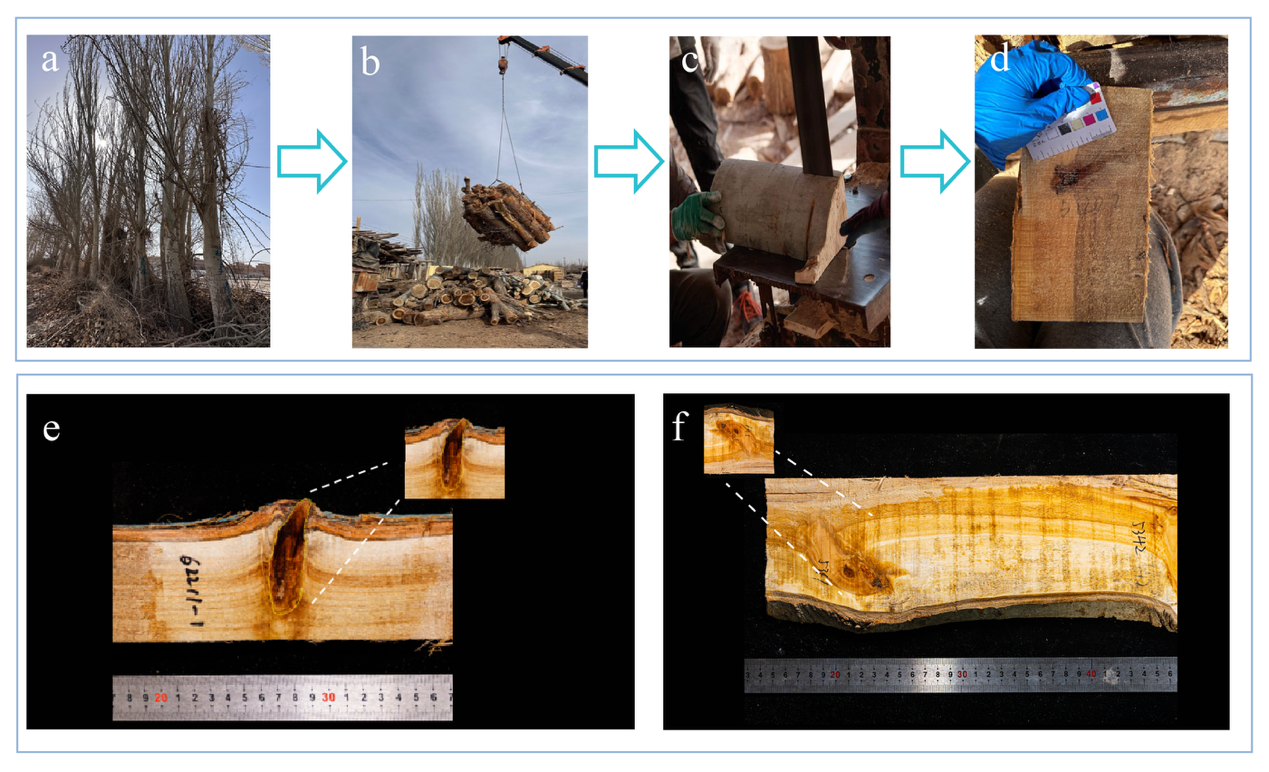

(a,b) Field sampling and collection of felled log materials. (c,d) Preparation of knot containing plank samples. (e,f) Representative validation images from *Populus euphratica* (e) and *Populus alba* var. *pyramidalis* (f), which were used for comparison between BioIMA and ImageJ measurements. Images include in frame scale references for calibration.

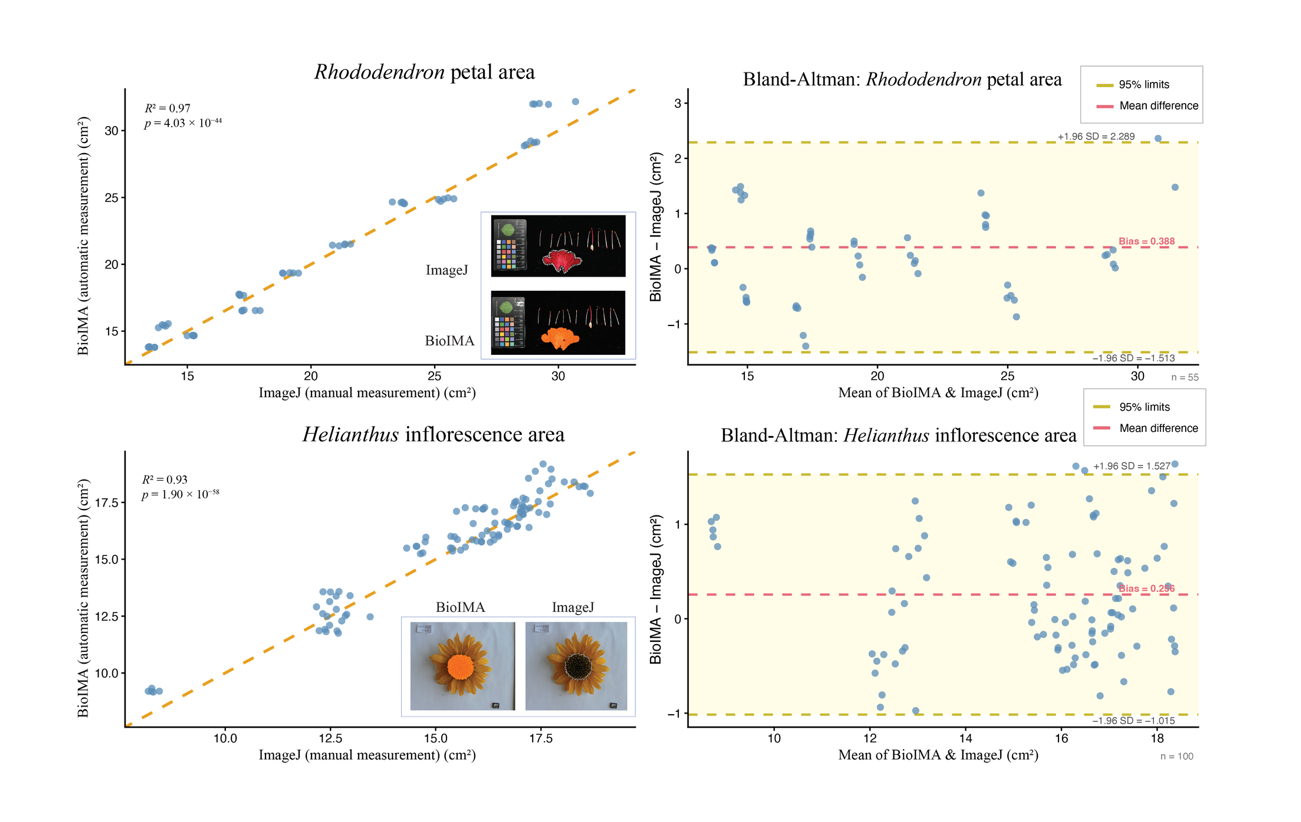

**Figure S2 | Additional validation of BioIMA area measurements on *Helianthus* inflorescence and *Rhododendron petal* images.**

BioIMA area measurements were compared with ImageJ manual measurements using independent *Rhododendron* petal and *Helianthus* inflorescence image datasets. The left panels show BioIMA-ImageJ measurement agreement, with orange dashed lines indicating the 1:1 identity line; *R*² and *p* values are shown in each panel. The right panels show Bland-Altman analyses, with red dashed lines indicating mean bias and yellow dashed lines indicating the 95% limits of agreement. Both datasets showed high agreement between methods.

**Table S1 | BioIMA exported trait fields and definitions.**

| **Category** | **Exported field** | **Description** | **Unit / format** |
| --- | --- | --- | --- |
| Metadata | Id | Unique object or measurement identifier | Integer or text |
| Metadata | Image Path | Path or file name of the source image | Text |
| Metadata | Label Name | User defined label for the selected region | Text |
| Metadata | Shape Type | Annotation type, such as polygon, line, or angle | Text |
| Morphology | Area | Calibrated area of the segmented region | User defined unit² |
| Morphology | Arap | Area in pixels | Pixels |
| Morphology | Perimeter | Calibrated boundary length | User defined unit |
| Morphology | Width | Width of the bounding box or selected region | User defined unit |
| Morphology | Height | Height of the bounding box or selected region | User defined unit |
| Morphology | Aspect Ratio | Width to height ratio | Unitless |
| Morphology | Circularity | Shape compactness derived from area and perimeter | Unitless |
| Morphology | Equivalent  Diameter | Diameter of a circle with the same area | User defined unit |
| Line and angle | Line Length | Calibrated length of a user defined line | User defined unit |
| Line and angle | Line Px | Line length in pixels | Pixels |
| Line and angle | Angle Degree | Angle measurement | Degree |
| Color | Mean RGB | Mean RGB value of the selected region | Text |
| Color | Hex Color | Hexadecimal Color code | HEX code |
| Color | Lab | Mean CIE Lab value of the selected region | Text |
| Color | MeanR | Mean red channel value | Numeric |
| Color | MeanG | Mean green channel value | Numeric |
| Color | MeanB | Mean blue channel value | Numeric |
| Color | LabL | Mean L value in CIE Lab space | Numeric |
| Color | LabA | Mean a value in CIE Lab space | Numeric |
| Color | LabB | Mean b value in CIE Lab space | Numeric |
| Color | Cluster K | Number of color clusters used for k-means color distribution analysis. | Integer |
| Color | Dominant Color Hex | HEX color code of the dominant color cluster within the selected region or mask. | HEX code |
| Color | Distribution | Relative proportions of the color clusters detected within ROI. | Proportion |

**Table S2 | Raw *Populus* knot measurement and timing benchmark data.** Raw BioIMA and ImageJ benchmark data for *Populus* knot images. The first sheet contains measurement time records used for the software throughput comparison in Fig. 3g. The second sheet contains paired morphometric measurements used for the method comparison and Bland-Altman analyses in Fig. 3c-f, including knot perimeter, major axis length, and knot area. (excel)

**Table S3 | Raw *Helianthus* inflorescence and *Rhododendron* petal validation data.** Raw BioIMA and ImageJ measurements for *Helianthus* inflorescence and *Rhododendron* petal area validation, including sample identifiers and measurement values used to generate Fig. S2. (excel)
