## Supplementary material for "BioIMA: a one-click desktop tool for standardized extraction of phenotypic traits from biological images": BioIMA User Manual (English)

### 1. Software Introduction

##### 1.1 Background

The observation, description, and quantification of external morphological traits are important for understanding similarities and differences among biological organisms. Quantifying variation in biological traits provides a foundation for a wide range of research fields. By standardizing the measurement of traits such as area, length, angle, color, and shape, researchers can compare individuals, populations, or species and further conduct statistical analyses.

However, because biological samples often differ greatly across species, many existing methods are designed for specific tasks and are not easily adapted or extended to new questions, image backgrounds, or datasets. In addition, some measurement tools lack sufficient standardization and reproducibility, which may introduce errors into downstream statistical analyses.

Traditional image-based measurement methods often rely on manual tracing, manual parameter adjustment, or semi-automated image-processing workflows. For example, when using software such as ImageJ, users may need to manually adjust thresholds, mark regions of interest, and extract phenotypic parameters step by step. Although these methods provide some flexibility, the workflow can be time-consuming and labor-intensive. The results may also be affected by user experience, image background, and parameter settings. For images with complex boundaries, uneven backgrounds, or large sample sizes, manual measurement can reduce both efficiency and the standardization and reproducibility of the final results.

##### 1.2 Main Functions of BioIMA

BioIMA is an intelligent recognition and automatic measurement tool designed for biological images. It aims to provide users with a more efficient, intuitive, and reproducible workflow for image-based phenotypic analysis. The software supports manual annotation and measurement, SAM-based segmentation and measurement, scale calibration, morphological measurement, color analysis, result preview, and CSV export.

Through simple interactive operations, users can select a target region, generate or edit a label, and automatically extract the corresponding phenotypic data.

##### 1.3 Basic Workflow

The main functions of BioIMA can be divided into three parts: the control module, the manual measurement module, and the automatic segmentation and measurement module.

The control module is used for image import, file management, result saving, and data export. The manual measurement module provides tools such as polygon, rectangle, line, angle, ruler, and color analysis. It is suitable for users who wish to manually select regions of interest and measure them. The automatic segmentation and measurement module is based on the SAM model. It allows users to quickly generate a mask for the target region using point prompts or box prompts, and then convert the mask into an editable and measurable label. In automatic mode, users only need to load the model, open an image, and provide a small amount of prompt information for the target region. The software can then generate a segmentation result automatically. After accepting the mask, BioIMA saves it as a label. Users can continue to inspect, edit, measure, and export the result. This workflow reduces repetitive manual tracing and improves the efficiency of measuring target regions in complex images.

BioIMA can be used for phenotypic measurement of plant organs, seeds, petals, leaves, wood structures, patterned regions, and other biological image samples. The software outputs measurements such as area, perimeter, width, height, aspect ratio, circularity, equivalent diameter, line length, angle, mean color values, and color composition ratios. These results can be directly used for downstream statistical analysis.

#### 2. Software Installation and Startup

##### 2.1 System Requirements

BioIMA is a cross-platform desktop application developed with Avalonia and .NET. It supports both Windows and macOS systems. Users can download the appropriate version according to their operating system. The recommended operating environment is as follows: Operating system: Windows 10 / Windows 11; macOS 12 or later. Hardware: At least 8 GB of memory is recommended. For high-resolution image processing, 16 GB of memory or higher is recommended for better performance. Storage: At least 1 GB of available storage space is recommended for the software, model files, image data, and analysis results. Image formats: BioIMA supports common image formats, including PNG, JPG, JPEG, BMP, and TIFF.

##### 2.2 Software Installation

BioIMA provides program versions for both Windows and macOS. Users should download the software package that matches their operating system. For Windows users, after downloading the software, users can either follow the installation wizard to complete the installation or open the extracted program folder and run the BioIMA executable file directly. For macOS users, after downloading the software, users can drag the BioIMA application into the applications folder, or launch the software directly from the downloaded or extracted folder. When running the software for the first time, macOS may ask users to confirm that they want to open an application from an identified or unverified developer. In this case, users should follow the system prompts to allow the application to open. During use, it is recommended not to move or delete files in the software installation directory. BioIMA may include built-in models, dependency libraries, and configuration files. Moving or deleting these files may cause some functions to stop working properly. It is also recommended to keep the software program and the data to be analyzed in separate locations. For example, the BioIMA application can be stored in the Applications folder or another fixed software directory, while the original images, analysis results, and exported files can be stored separately in a project folder.

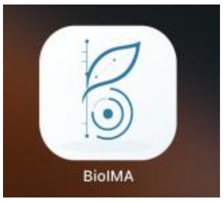

##### 2.3 Starting the Program

After installation, users can start BioIMA by double clicking the program icon. Once launched, the software will open the main interface, where users can import images, create annotations, perform SAM-based segmentation, set the scale, measure phenotypic traits, and export results.

If no image is displayed after the software starts, it means that no image has been imported yet. Users should first use the **Open Image** or **Open Folder** function to select the image files to be analyzed.

If users need to use the SAM-based segmentation function, the software will enter the segmentation-ready state after the model has been loaded. The current version includes a built-in model, so users can usually use this function directly without manually loading an additional model file.

##### 2.4 Workspace and File Management

When using BioIMA for image analysis, it is recommended that users create a separate workspace folder for each project. This folder can be used to store the original images, analysis results, and exported files. A clear file structure helps with later checking, repeated analysis, and data organization. A recommended project folder structure is shown below:

```
Project_Name/
├──── images/          stores the original image files
├──── results/         stores exported CSV measurement results
├──── color_reports/ stores exported color analysis reports in PNG format
└──── models/          stores custom model files, optional
```

BioIMA already includes a built-in SAM model, so users usually do not need to create a separate model’s folder. A separate model folder is only recommended when users need to use custom models or manage different versions of model files.

To ensure that results are traceable, users are advised not to directly modify the original image files. The original images can be stored in the images folder, measurement results can be exported to the results folder, and color analysis reports can be saved in the color reports folder.

This structure clearly separates raw data from analysis outputs and makes downstream statistical analysis and result checking easier.

### 3. Software Interface Overview

#### 3.1 Overall Layout of the Main Interface

The main interface of BioIMA consists of several functional areas, including the left toolbar, the central image display area, the Labels panel, the Mask Preview panel, and the Results panel. Users can complete image import, annotation creation, SAM-based segmentation, result preview, phenotypic measurement, and data export within the same interface. After launching the software, the main interface is shown as follows:

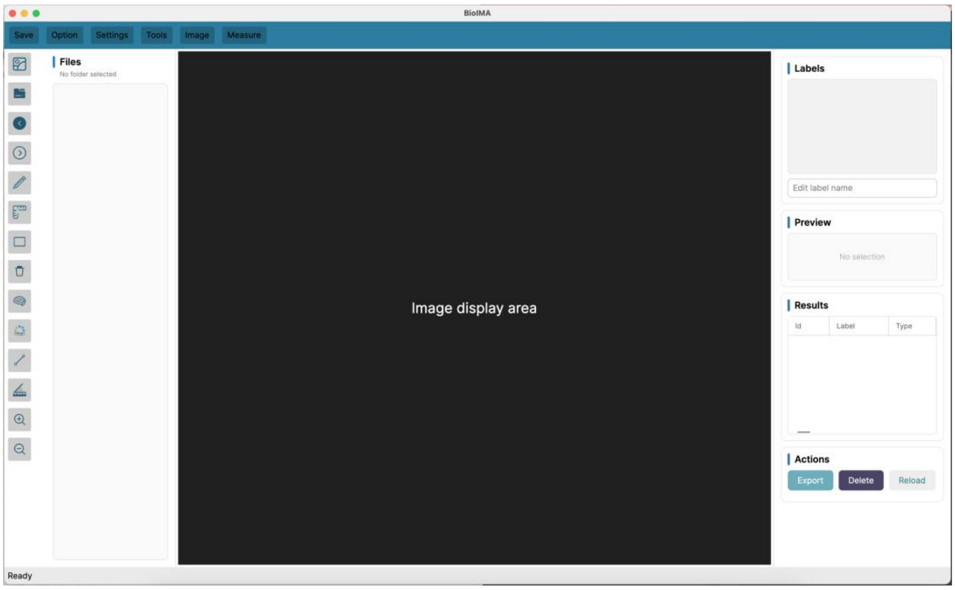

Click the Open Image icon to access the user's local folders. The interface is shown as follows:

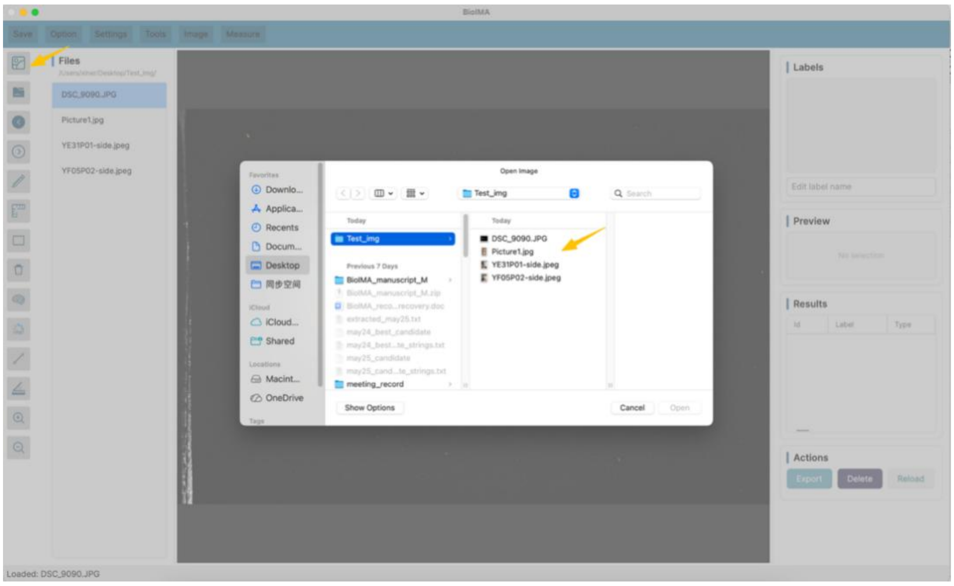

##### 3.2 Left Toolbar and Image File List

The left side of the BioIMA interface contains the main operation tools and the image file list. The vertical toolbar is used for common operations, such as opening images, creating annotations, setting the scale, performing measurements, and zooming in or out. The image file list displays the images in the currently opened folder, allowing users to quickly switch between different samples.

The left toolbar includes the following functions:

- Open Image 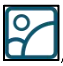/ Open Folder 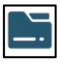: Open a single image or an image folder.
- 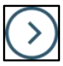 Switch to the next image in the current folder.
- 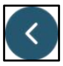 Switch to the previous image in the current folder.
- 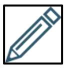 Draw a polygonal region, suitable for measuring targets with irregular boundaries.
- 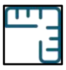 Set the image scale, allowing pixel-based measurements to be converted into real-world units.
- 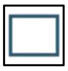 Draw a rectangular region, suitable for quickly selecting regular areas.
- 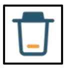 Draw a rectangular area to quickly remove edited content within the selected region.
- 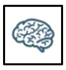 Start the SAM-based segmentation function and generate a mask using point prompts or box prompts.

- 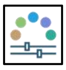 Analyze the selected region and calculate the mean color and color composition ratio.

  *Note: Color analysis is also performed automatically after clicking Measure for manual or SAM-generated annotations.*
- 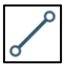 Draw a line segment for length measurement.
- 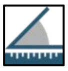 Draw an angle for angle measurement.
- 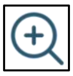 Enlarge the image view.
- 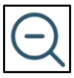 Reduce the image view.

Image File List:

The Files image file list is located to the right of the vertical toolbar. It is used to display the image files in the currently opened folder. After a user opens an image folder, BioIMA lists the images in the Files panel. Users can click a file name in the list to switch the image currently being analyzed. The main functions of the image file list include: displaying the image files in the current folder; quickly switching between different sample images; helping users confirm the name of the image currently being analyzed; working together with the Results table to record image paths and sample information.

##### 3.3 Top Toolbar and Image List

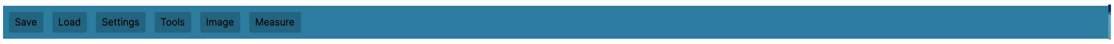

The top toolbar provides access to the main workflow of the software, including saving project data, loading images or annotation files, adjusting default settings, selecting annotation and analysis tools, performing image processing operations, and carrying out measurements.

###### 3.3.1 Save

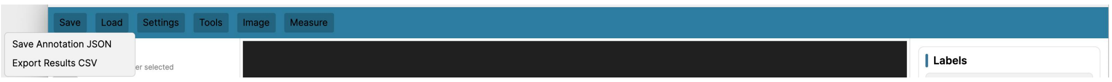

Use **Save Annotation JSON** to save the current annotation project as a JSON file. The saved file includes label information, shape types, annotation coordinates, label names, label colors, and accepted SAM mask-backed labels.

This function is useful when users want to continue editing or measuring the same image later.

**Steps**

1.Open or annotate an image.

2.Click **Save** in the top toolbar.

3.Select **Save Annotation JSON**.

4.Choose a save location and file name.

5.BioIMA saves the current annotation data as a JSON file.

**Note**: The annotation JSON file stores annotation information, not the original image itself. Users should keep the original image file together with the annotation JSON file.

Use **Export Results CSV** to export the current measurement table as a CSV file. The exported table may include morphological measurements, color measurements, color distribution results, and the selected color cluster K value.

This function is the same as the **Export** button in the Actions panel. (This function works the same way as the Export button in the Actions panel on the right) .

**Steps**

1.Measure one or more labels.

2.Click **Save** in the top toolbar.

3.Select **Export Results CSV**.

4.Choose a save location and file name.

5.BioIMA exports the Results table as a CSV file.

##### 3.3.2 Load

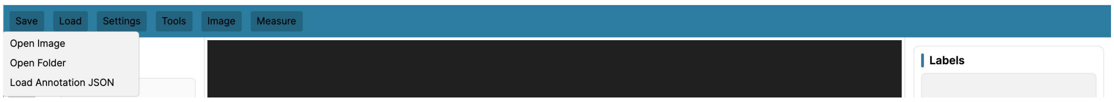

The “**Load**” menu contains functions for opening images, opening image folders, and loading previously saved annotation files.

Use **Open Image** to open a single image for subsequent annotation, segmentation, and measurement. Supported image formats include PNG, JPG, JPEG, BMP, TIFF, as well as HEIC/HEIF files if supported by the current environment.

Use **Open Folder** to load a folder containing multiple images. The image list will be displayed in the Files panel on the left, allowing users to switch between different images during analysis.

Use **Load Annotation JSON** to load a previously saved BioIMA annotation file. This function restores the saved labels, shapes, coordinates, colors, and accepted SAM mask labels.

**Steps**

1. Open the corresponding original image.

2. Click Load in the top toolbar.

3. Select Load Annotation JSON.

4. Choose the previously saved annotation JSON file.

5. The saved annotation content will be restored.

**Note**: To ensure coordinate accuracy, it is recommended to load the annotation JSON file using the same image that was used when the annotations were saved.

##### 3.3.3 Settings

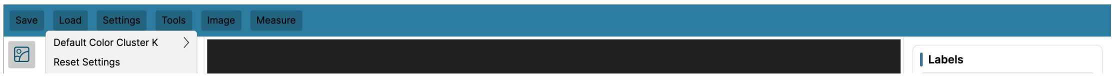

Use **Default Color Cluster K** to set the default number of color classes used in color distribution analysis. The K value determines how many color clusters BioIMA will identify within a selected region or mask.

For example, **K = 5** means that BioIMA will divide the selected pixels into five representative color classes.

**Steps**

1.Click Settings in the top toolbar.

2.Select Default Color Cluster K.

3.Choose a K value.

4.The selected K value will be used as the default for subsequent color analysis.

##### 3.3.4 Tools

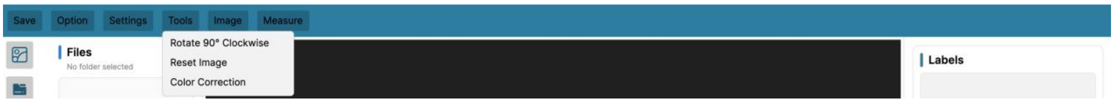

The "**Tools**" menu allows users to select annotation, measurement, segmentation, and color analysis tools.

Use **Polygon** to manually outline the target contour by placing multiple points. This tool is suitable for annotating irregular structures.

**Steps**

1.Click Tools in the top toolbar.

2.Select Polygon.

3.Click along the target boundary to add points.

4.Double-click to complete the polygon.

Use **Rectangle** to annotate a rectangular region by selecting two opposite corners.

**Steps**

1.Click Tools in the top toolbar.

2.Select Rectangle.

3.Click the first corner of the rectangular region.

4.Click the opposite corner to complete the rectangle annotation.

Use **Line** to measure the distance between two points.

**Steps**

1.Click Tools in the top toolbar.

2.Select Line.

3.Click the starting point of the line.

4.Click the ending point of the line.

5.Record the line length.

Use **Angle** to measure the angle formed by three points.

**Steps**

1.Click Tools in the top toolbar.

2.Select Angle.

3.Click three points in sequence to define the angle.

4.Calculate the angle value.

Use **Ruler** to set the image scale using a known length. After calibration, pixel-based measurements can be converted into real-world units.

**Steps**

1.Click Tools in the top toolbar.

2.Select Ruler.

3.Click the two endpoints along the known length.

4.Enter the real length and unit.

5.Apply the scale to subsequent measurements.

Use **Segmentation** to generate a target mask with assistance from the SAM model. Users can guide the segmentation using positive/negative point prompts or a box prompt, and then accept the generated mask as an editable label.

**Steps**

1.Click Tools in the top toolbar.

2.Select SAM Segmentation.

3.Use positive points, negative points, or a box prompt to guide the segmentation.

4.Preview the generated mask.

5.Click Accept Mask to convert the mask into a label.

6.If needed, continue measuring the mask label.

Use **Color Box** to analyze the color within a rectangular region. The software calculates the mean RGB, Hex, and Lab values, as well as the color distribution of the selected region.

**Steps**

1.Click Tools in the top toolbar.

2.Select Color Box.

3.Click two opposite corners to define the rectangular region.

4.The software will open the Color Analysis window.

5.View the mean color and color distribution results.

6.Save the results if needed.

##### 3.3.5 Image

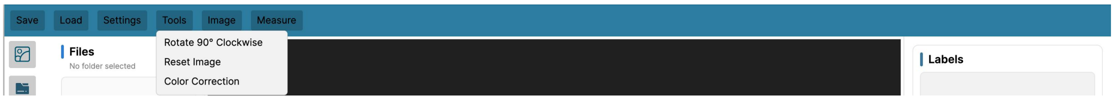

The Image menu provides basic image adjustment functions for processing the current image before annotation, segmentation, and measurement. In the current version, the Image menu mainly includes three functions: Rotate 90 ° Clockwise, Reset Image, and Color Correction. These functions are used to rotate the image, restore the original image state, and perform basic color correction, respectively.

**1. Rotate 90 ° Clockwise:**

Rotate 90 ° Clockwise is used to rotate the current image 90 ° clockwise. This function can be used when the image orientation is incorrect or when users want to annotate and measure the image in a more convenient orientation. After this function is applied, BioIMA will also update the coordinates of existing annotation objects, so that the previously drawn labels remain aligned with the rotated image as much as possible.

**Steps**

1.Open the image to be analyzed.

2.Click Image in the top menu bar.

3.Select Rotate 90 ° Clockwise.

4.The current image will be rotated 90 ° clockwise.

5.The positions of existing labels will be automatically updated according to the rotated image.

**Note**: If SAM segmentation has already been performed on the current image, the temporary SAM mask and image embedding will be cleared after the image is rotated. If users still need to use SAM-assisted segmentation, they should perform SAM segmentation again after rotating the image.

##### 2. Reset Image

Reset Image is used to reload the original image from disk. This function can be used when users have rotated the image or applied color correction, but want to restore the image to its original state when it was first opened.

**Steps**

1.Click Image in the top menu bar.

2.Select Reset Image.

3.BioIMA will reload the original image file.

4.The currently displayed image will be restored to the original version.

**Note**: In the current version, Reset Image will also clear the annotations, labels, temporary SAM mask, and measurement state on the current image. This is designed to avoid coordinate inconsistencies between the original image and the modified image. Therefore, before using Reset Image, users are advised to confirm whether the current results have already been saved or exported.

#### 3. Image Color Correction

The **Color Correction** function is used to reduce color bias caused by lighting conditions, camera settings, or scanning conditions. This function performs basic white balance or gray reference correction on the current image based on a white or gray reference area selected by the user.

Color correction is based on a neutral reference area selected by the user. This area should theoretically be close to white or gray in the image, such as a gray card, white paper, scale ruler, white scale bar, or a neutral color standard block. The software calculates the mean RGB value of the selected reference area and adjusts the RGB channels of the entire image based on this result, thereby reducing overall color bias.

##### 3.3.6 Measure

The **Measure** button is used to measure the currently selected label. Depending on the label type, BioIMA can calculate area, perimeter, width, height, aspect ratio, circularity, equivalent diameter, line length, angle, mean color, Lab color values, and color distribution.

##### Steps

1.Select a label in the Labels panel.

2.Click Measure in the top toolbar.

3.BioIMA will calculate the available measurement results for the selected label.

4.The results will be displayed in the Results table.

5.Users can export the results through Save → Export Results CSV or the Export button in the Actions panel on the right.

#### 3.4 Image Display Area

The image display area is in the center of the main interface and is used to display the currently opened image. Most interactive operations are performed in this area, such as drawing polygons, selecting rectangular regions, setting a ruler, adding prompt points, viewing SAM segmentation results, and editing label boundaries.

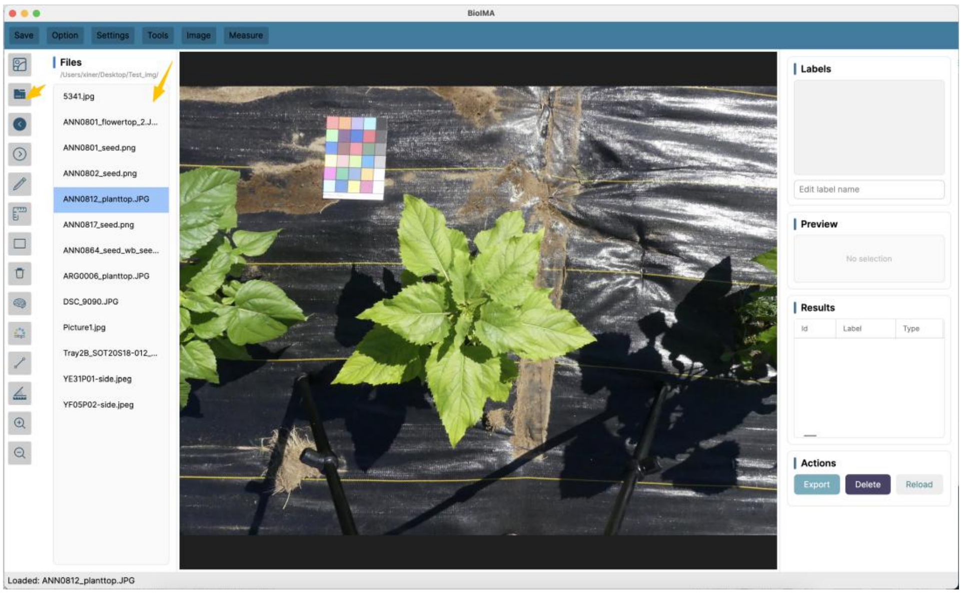

##### 3.5 Labels Panel

The Labels panel displays all labels created for the current image. Labels created manually, such as polygon and rectangle annotations, as well as labels generated by accepting a SAM mask, will all appear in this list. Users can use the Labels panel to: view all labels in the current image; select a label for measurement or editing; check the label name and display color; rename a label; delete labels that are no longer needed; switch between different labels for measurement.

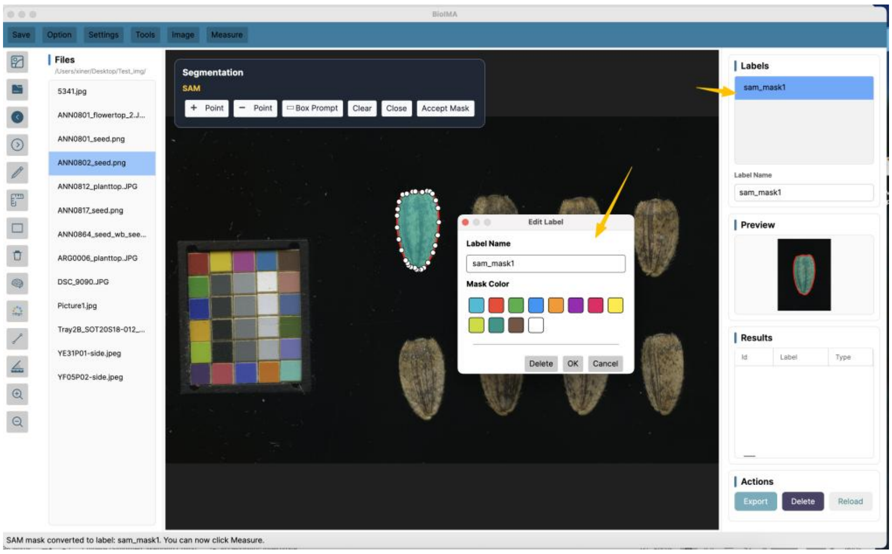

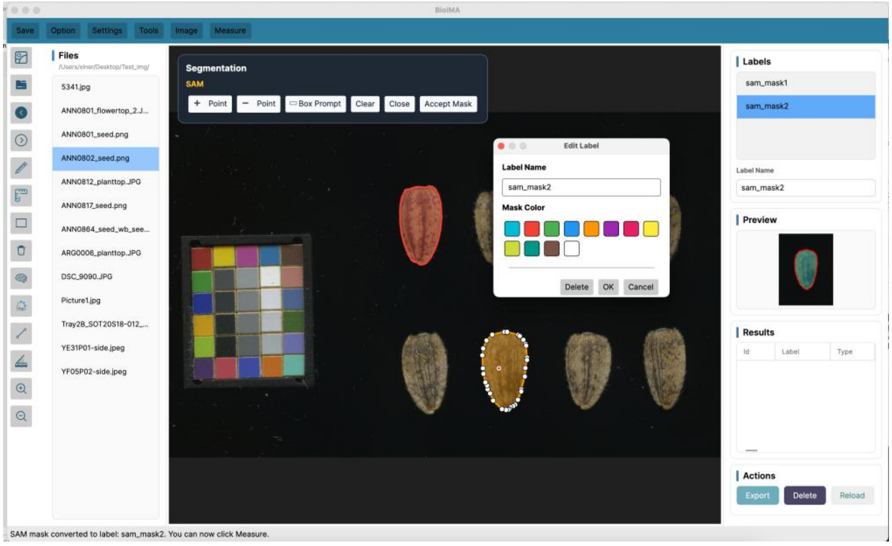

When a user selects a label, the corresponding boundary or mask location will be displayed in the image display area. The Mask Preview panel will also show a local preview of the selected label. Before measurement, users should make sure that the selected label matches the intended target region, so that incorrect regions are not saved to the results table.

##### 3.6 Mask Preview Panel

The Mask Preview panel shows a local preview of the currently selected label. The preview usually includes the local region of the original image, the label boundary, and the mask overlay. This helps users quickly check whether the selected region is correct. The Mask Preview panel can be used to: confirm whether the selected label corresponds to the target region; check whether a SAM mask fully covers the target; determine whether the boundary needs further editing; quickly verify the analysis region before measurement.

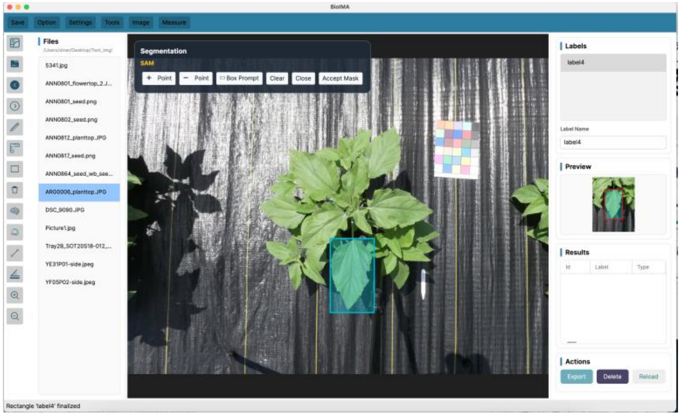

For manually drawn labels, the Mask Preview panel displays the outline of the selected region. For labels generated by SAM-based segmentation, it displays the local image region and the corresponding mask coverage. If the user has edited the boundary of a SAM-generated label, the preview can also be used to check the modified region.

If no label is selected, or if the selected label does not contain a region that can be previewed, the Mask Preview panel may be empty. Users can first select a label in the Labels panel and then view the corresponding preview.

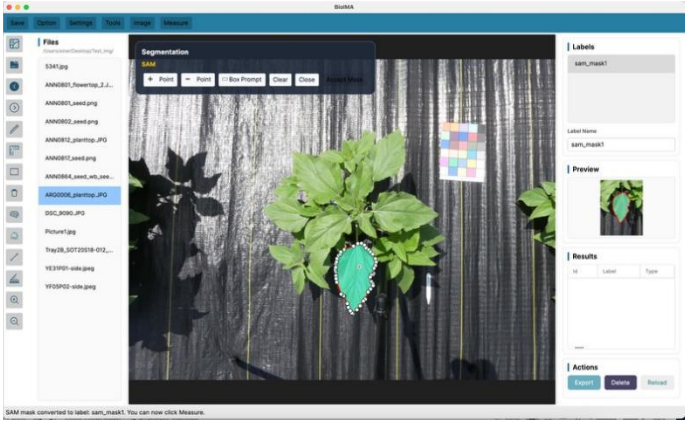

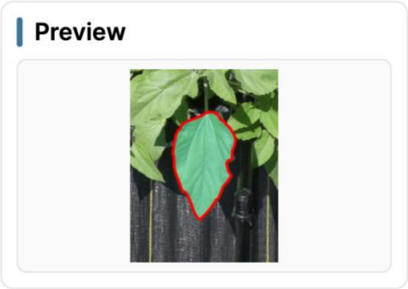

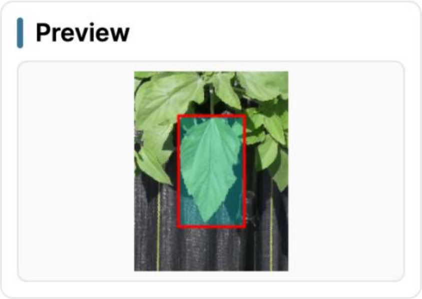

##### 3.7 Results Panel

The Results panel is used to display the measurement results that have been saved by the user. Each time a user measures and saves a label, line segment, angle, or color analysis region, a new record is added to the Results table. The Results panel may contain the following types of information: image path; label name; shape type; area and pixel area; perimeter, width, and height; aspect ratio, circularity, and equivalent diameter; line length and angle; mean RGB, Hex, and Lab color values; number of color classes, K; dominant color and color distribution.

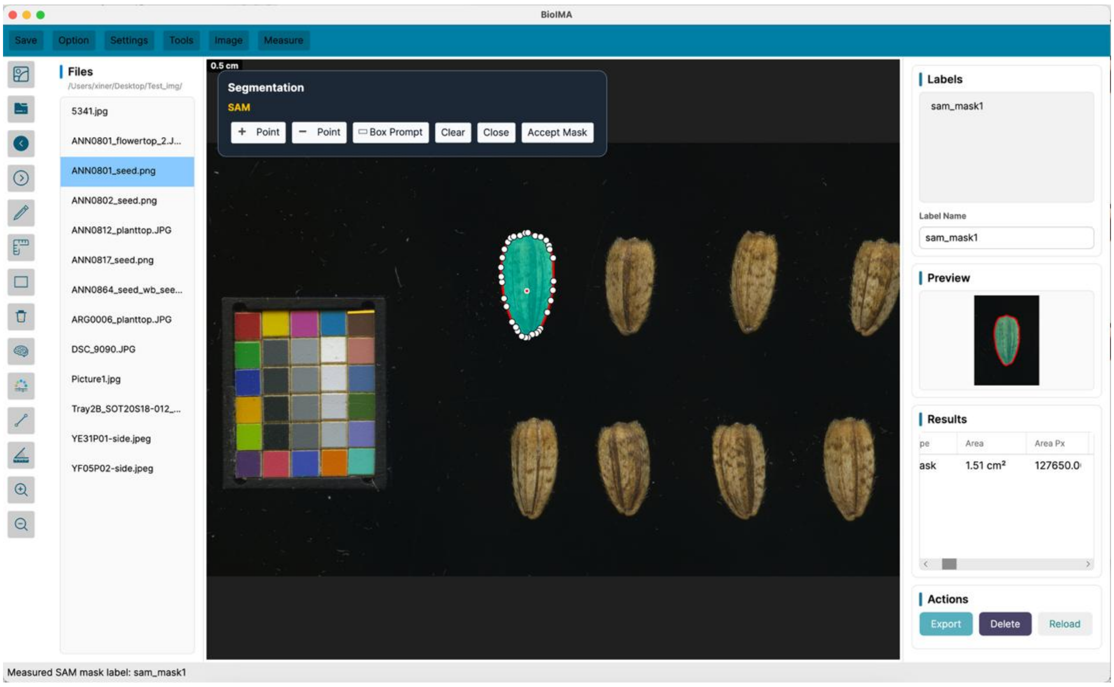

| Id | ImagePath | LabelName | ShapeType | Area | AreaPx | Perimeter | Width | Height | AspectRatio | Circularity | EquivalentDiameter |
|---|---|---|---|---|---|---|---|---|---|---|---|
| ANN0801_seed | /Users/xiner/Desktop/Test_img/ANN0801_seed.png | sam_mask1 | mask | 1.51 cm² | 127650.00 | 5.92 cm | 0.99 cm | 1.97 cm | 0.5009 | 0.5422 | 1.39 cm |

| LineLength | LinePx | AngleDegree | MeanRgb | HexColor | Lab | MeanR | MeanG | MeanB | LabL | LabA | LabB | ColorClusterK | DominantColorHex | ColorDistribution |
|---|---|---|---|---|---|---|---|---|---|---|---|---|---|---|
| NA | NA | NA | 120, 102, 66 | #786642 | 44.02, 2.04, 22.73 | 120.11 | 101.76 | 66.33 | 44.02 | 2.04 | 22.73 | NA | #675436 | #675436:25.1%; #836C41:23.5%; #968658:20.4%; #7A6B4B:20.1%; #4D3C2B:10.9% |

Users can check the saved measurement records in the Results panel and export the results as a CSV file after completing the analysis. The exported CSV file can be used for downstream statistical analysis, plotting, or integration with other phenotypic, genotypic, or environmental data. Before exporting the results, users are advised to check each record in the Results table, including the image name, label name, and measurement fields, to ensure that the results correspond to the correct samples.

### 4. Image Import and Basic Operations

BioIMA supports both single-image import and folder-based image import. Users can choose the appropriate import method according to their analysis needs. For single-image analysis, users can open the target image directly. For batch analysis, it is recommended to place images from the same batch in one folder and use the folder import function to switch between images and measure them one by one.

#### 4.1 Opening a Single Image

Users can import a single image by clicking the Open Image button in the menu bar or the left toolbar. After clicking the button, a file selection window will appear, allowing users to select the image to be analyzed from a local folder.

#### 4.2 Opening an Image Folder

When users need to analyze a batch of sample images, they can use the Open Folder function. After a folder is opened, BioIMA reads the image files in that folder and displays the file names in the Files image list on the left side. Users can click different file names in the list to quickly switch the current image.

#### 4.3 Image Switching and Display

After an image folder is opened, the Files list on the left displays the image files in the current folder. Users can switch the current image by clicking a file name. The selected image will be displayed in the central image display area. When switching images, users should make sure that the current image matches the sample they intend to analyze. Before annotation or measurement, it is recommended to check the selected file name in the Files list and the current image name shown in the status bar at the bottom of the interface. The image display area is used to view the original image and its related annotations. Depending on the current operation, the image may show manually drawn polygons, rectangles, line segments, or angles; ruler lines; SAM-generated mask overlays; saved label boundaries; and the highlighted state of the currently selected label. If the image display scale is not suitable, users can use the Zoom In or Zoom Out tools to adjust the view. Zooming only changes how the image is displayed in the interface. It does not modify the original image file and does not affect the measurement results.

##### 4.4 Undo, Redo, and Clearing Operations

During image analysis, users may need to modify or remove recently created annotations or temporary results. BioIMA provides several basic editing and clearing functions to help users adjust annotations or reselect analysis regions. Common operations include:

1. Users can use the rectangular erase tool 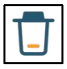 to quickly remove annotation content within a selected rectangular area. After clicking the erase icon, users can define a rectangle on the image by clicking two points. All annotation content within the rectangle, such as angles, line segments, and polygons, will be removed.

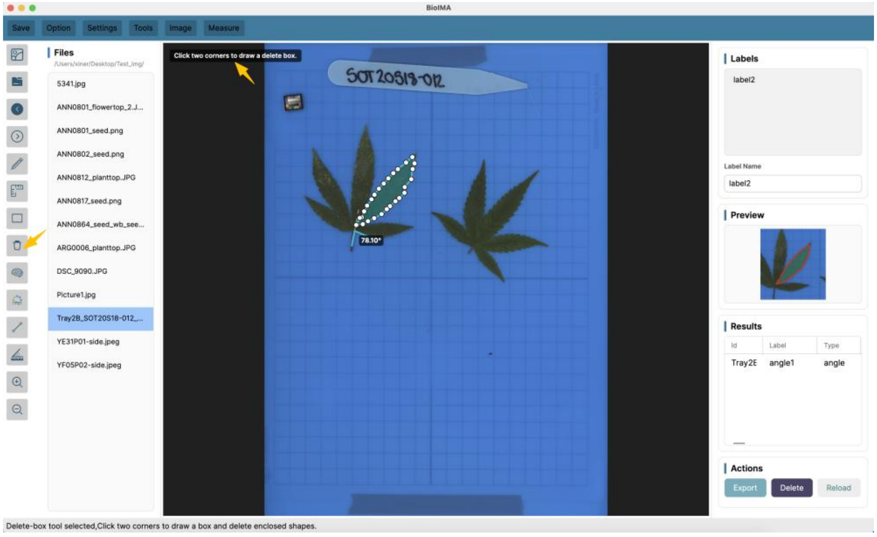

2. Users can click the Delete button to delete the label object that is currently being previewed or selected.

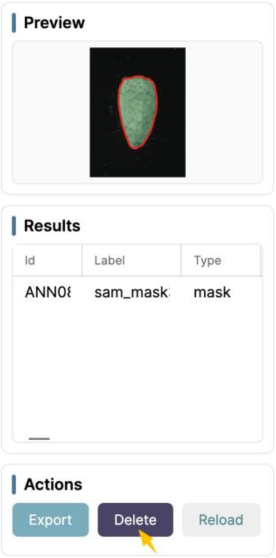

3. Users can click the Reload button to remove all objects currently displayed on the panel.
4. Users can double-click a label name to rename it. They can also use the delete button in the Edit Label panel to remove the selected object.

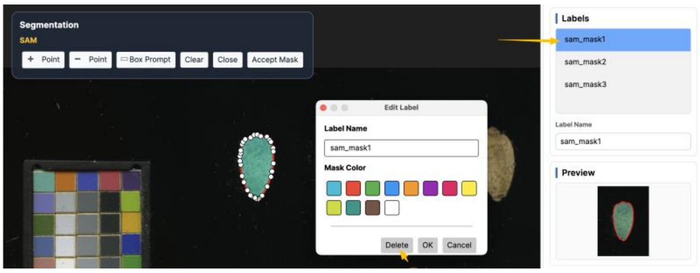

In the segmentation panel, the Clear button is used to remove the current prompt points, prompt box, or temporary segmentation result, allowing users to reselect the target region. If the user has already clicked Accept Mask and saved the mask as a label, the result will appear in the Labels panel. In this case, to remove the label, users should select it in the Labels panel and then use the delete function.

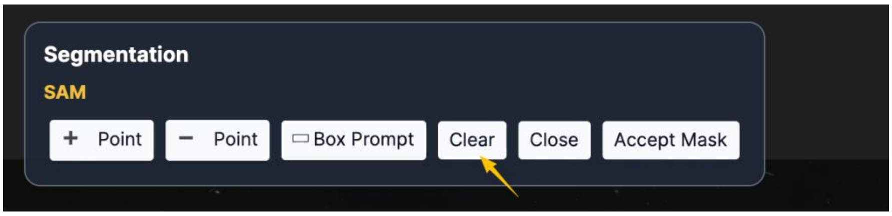

Before deleting a label or a measurement result, users should confirm that the currently selected object is correct to avoid accidentally removing completed annotations or saved records. For important results, it is recommended to export the CSV file in time for backup. BioIMA supports the full workflow of image import, manual annotation, SAM mask conversion to label, measurement result saving, and CSV export.

### 5. Scale Calibration

When measuring biological images, the software records measurements in pixel-based units by default, such as pixel area and pixel length. If users need area or length measurements in real-world units, scale calibration must be performed first. BioIMA provides a ruler-setting function. Users can select a line segment in the image with a known real-world length and enter the corresponding length and unit. After calibration, the software converts subsequent measurements into real-world units based on the relationship between pixel length and actual length. Scale calibration is usually needed in the following situations: the image contains a scale bar; the sample was photographed or scanned at a fixed resolution; the user knows the actual length of a structure in the image; the user needs to compare real-world area or length across different images. If users only need relative comparisons or pixel-based results, scale calibration is not required. In this case, the software can still output pixel area and pixel length.。

##### 5.1 Setting the Ruler Line

Before setting the scale, users should identify a region in the image with a known real-world length. This may be a scale bar included in the image, a ruler mark in a scanned image, or a reference object with a known size. The operation steps are as follows: 1. Open the image to be analyzed. 2. Select the Ruler tool from the left toolbar. 3. Click the starting point of the ruler line in the image display area. 4. Move the cursor to the endpoint of the ruler line and click again to complete the line. 5. The software will calculate the pixel length of the line drawn by the user.

When drawing the ruler line, users should click as accurately as possible on the two ends of the region with known length. If the image contains a scale bar, it is recommended to align the ruler line exactly with the scale bar. The more accurately the ruler line is drawn, the more reliable the subsequent area and length conversions will be.

If the ruler line is not placed accurately, users can redraw it or reset the scale and set it again.

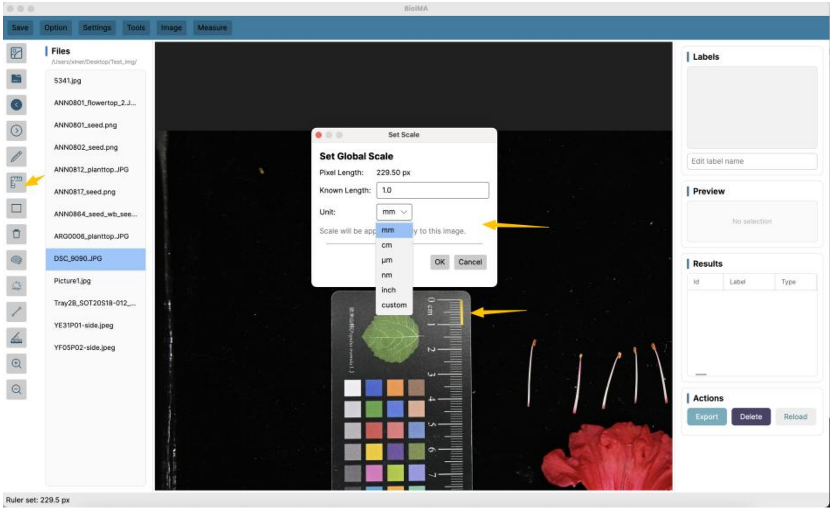

##### 5.2 Entering the Real-World Length and Unit

After drawing the ruler line, users need to enter the real-world length and unit corresponding to that line segment. The real-world length should match the actual length of the selected segment in the image. After the length and unit are entered, the software establishes the conversion relationship between pixels and real-world units. When users later measure area, length, or line segments, BioIMA can output both pixel-based results and real-world measurements. Users should be careful not to confuse different units when entering the length. If the unit is entered incorrectly, all subsequent measurements in real-world units will be affected.

##### 5.3 Conversion Between Pixel Units and Real-World Units

The key purpose of scale calibration is to establish the relationship between pixel length and real-world length. Based on the pixel length of the ruler line drawn by the user and the real-world length entered by the user, the software calculates the real-world distance represented by each pixel. The length conversion is calculated as: Real-world length = Pixel length × Real-world length per pixel. The area conversion is calculated as: Real-world area = Pixel area × (Real-world length per pixel)$^2$. Therefore, after scale calibration, BioIMA can save both pixel area and real-world area. Pixel area is suitable for relative comparisons within images, while real-world area is more suitable for standardized comparisons across different images, samples, or experiments.

##### 5.4 Resetting the Scale

If users find that the scale was set incorrectly, or if they need to set a new scale for another image, they can use the reset scale function. After resetting, the current conversion relationship between pixels and real-world units will be cleared. Users can then draw a new ruler line and enter a new real-world length. After the scale is reset, and before a new calibration is completed, the software can only output measurements in pixel units. If real-world area or real-world length is needed, users should set the ruler line again and enter the correct length and unit. For a batch of images captured with the same scanning parameters or imaging scale, the same scale setting may be used. However, if different images have different resolutions, zoom levels, or shooting distances, scale calibration should be performed separately for each image to ensure accurate measurements.

#### 6. Manual Annotation and Measurement

BioIMA supports manual annotation. Users can choose the appropriate tool according to the shape of the target region and the purpose of measurement. For irregular regions, the polygon tool can be used. For regular regions, the rectangle tool is suitable. For distance or length measurements, users can use the line tool. For angle measurements, the angle tool can be used.

Manual annotation is suitable for images where the target boundary is clear and where users want to define the analysis region themselves. After an annotation is created, the software saves the region as a label and allows users to further edit, measure, and export the results. Manual annotations can be displayed together with SAM-basedsegmentation results in the Labels panel and saved in the Results table. BioIMA currently supports manual tools such as polygon, rectangle, line, angle, and ruler, and the measurement results can be exported as a CSV file.

##### 6.1 Polygon Region Annotation

Polygon annotation is suitable for target regions with irregular boundaries, such as tree knots, leaves, petals, seeds, patterned regions, or other biological structures with complex shapes. Users can click multiple points along the target boundary to create a closed polygon region.

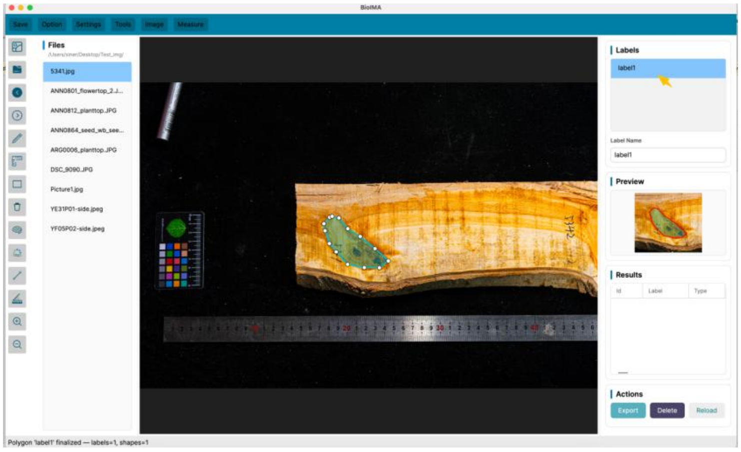

The operation steps are as follows: 1. Open the image to be analyzed. 2. Select the Polygon tool from the left toolbar. 3. Click along the target boundary in the image display area to add polygon vertices. 4. After selecting the target region, finish the drawing to generate a label. 5. Check the newly generated label in the Labels panel on the right. 6. To measure the region, select the label and click Measure.。

When drawing a polygon, the vertices should be placed as closely as possible along the target boundary. For regions with complex boundaries, users can add more vertices to improve the fit of the outline. For regions with simpler boundaries, fewer vertices are usually sufficient and can make later editing easier. After the polygon annotation is completed, users can view its boundary in the image display area or check the local preview of the label in the Mask Preview panel. If the boundary is not accurate, it can be adjusted using the vertex editing function.

##### 6.2 Rectangle Region Annotation

Rectangle annotation is suitable for regular regions or for quickly selecting a local area of an image. For example, users can use the rectangle tool to select a fixed sample region for later color analysis, local measurement, or quick recording.

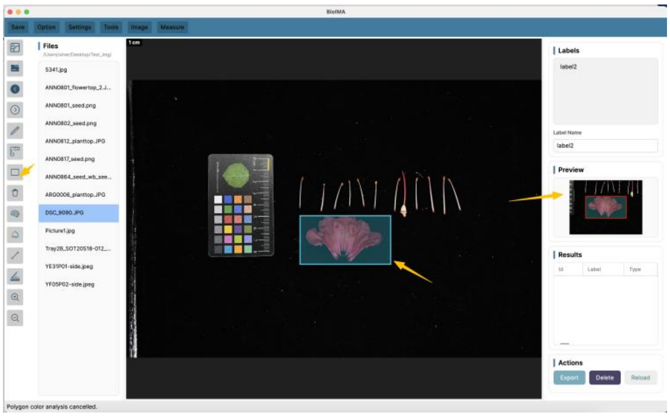

The operation steps are as follows: 1. Select the Rectangle tool from the left toolbar. 2. Click and drag in the image display area to select the target region. 3. Release the mouse to generate a rectangular annotation. 4. Check the newly created label in the Labels panel. 5. Select the label and click Measure to save the measurement result.

The rectangle tool is suitable when the target has a regular shape or when users only need to analyze a fixed local region. If the target boundary is irregular, it is recommended to use the Polygon tool or the SAM-basedsegmentation function to obtain a region that better follows the target boundary. After a rectangle annotation is created, it will also appear as a label in the Labels panel. Users can select the label, check its position in the image, and measure or delete it as needed.

##### 6.3 Line Length Measurement

The Line tool is used to measure the distance between two points. It is suitable for measuring sample length, width, the distance between structures, or other linear traits. If scale calibration has been completed, the software can convert pixel length into real-world length. The operation steps are as follows: 1. Select the Line tool from the left toolbar. 2. Click the starting point of the line segment in the image. 3. Move to the endpoint and click again to complete the line. 4. Click Measure or save the measurement result. 5. View the line length in the Results panel.

If no scale has been set, the software will output the line length in pixel units. If scale calibration has been completed, the Results panel can also save the length in real-world units. When using the Line tool, users should select the starting and ending points as accurately as possible. For samples that require repeated measurements, it is recommended to use a consistent measurement rule. For example, users may always measure from the leftmost point to the rightmost point of a structure, or always measure along the main axis, to ensure that the results are comparable.

##### 6.4 Angle Measurement

The Angle tool is used to measure the angle between two line segments. It is suitable for analyzing leaf opening angles, branch angles, organ bending angles, or other geometric traits. The operation steps are as follows: 1. Select the Angle tool from the left toolbar. 2. Click three points in the image to define the angle. 3. In general, the first and third points are located on the two sides of the angle, and the second point is the vertex. 4. After the three points are selected, the software calculates the angle. 5. Click Measure or save the result, and view the angle value in the Results panel.

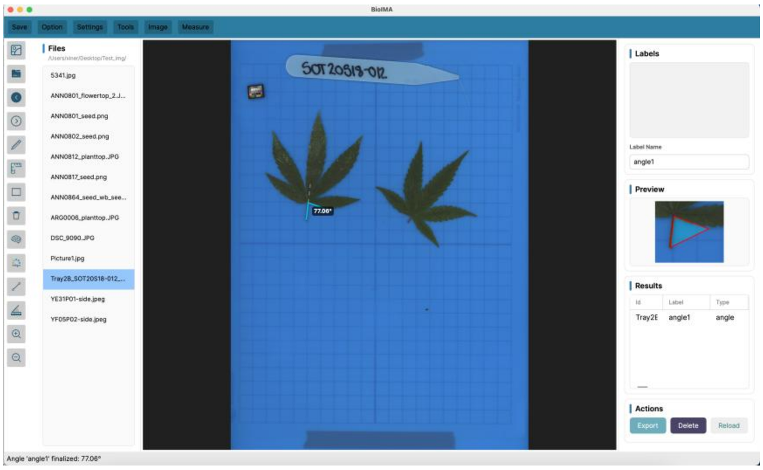

Angle measurements are usually reported in degrees. Before measurement, users should clearly define how the angle is measured and use the same point-selection rule for all images in the same batch. For example, when measuring leaf angle, users should always select the three points in the same order and at the same structural positions. If the selected points are not accurate, users can delete the current angle annotation and draw it again.

##### 6.5 Editing Annotation Vertices

For existing polygon, line, or angle annotations, users can adjust the annotation position using the vertex editing function. This function is useful when the boundary is not accurate enough after annotation, or when some vertices need minor adjustment. Common editing steps include: Select an existing label; View its vertices in the image display area; Drag a vertex to a new position; Release the mouse to complete the modification; Check the annotation boundary again in the image display area or in the Mask Preview panel.

For polygon regions, users can adjust the boundary shape by moving the vertices. For line and angle measurements, users can move the starting point, endpoint, or angle vertex to correct the measurement position.

If a label was generated from a SAM mask and the user manually edits its boundary, the software will use the edited boundary for subsequent measurements. If the boundary has not been edited, the SAM mask label will retain the measurement accuracy of the original mask by default.

##### 6.6 Measurement Results of Manual Annotations

After completing a manual annotation, users can select the corresponding label and click Measure to perform measurement. The results will be displayed in the Results panel and can later be exported as a CSV file.

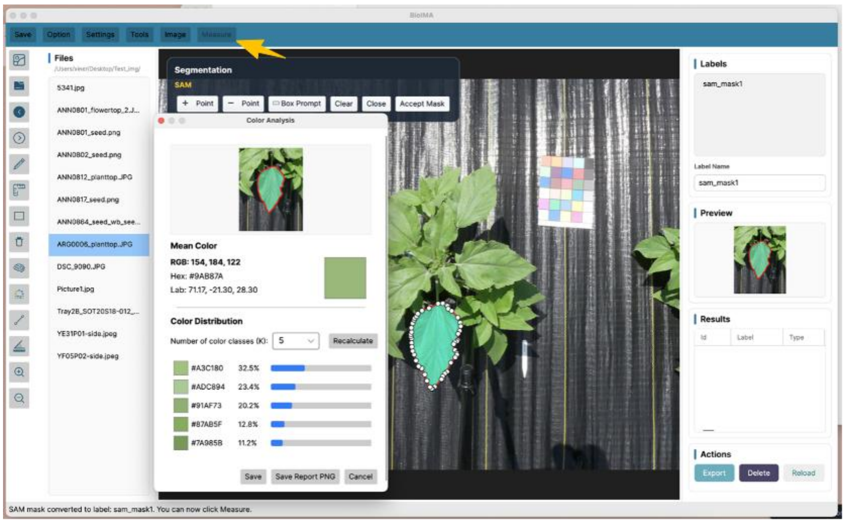

After completing a manual annotation, users can select the corresponding label and click Measure to perform measurement. The results will be displayed in the Results panel and can later be exported as a CSV file.

For region-based annotations, such as polygon and rectangle, the software usually outputs the following measurements: Area: the area of the region. If scale calibration has been completed, the area can be reported in real-world units; Area Px: the pixel area of the region; Perimeter: the perimeter of the region; Width: the width of the region; Height: the height of the region; Aspect Ratio: the ratio between width and height; Circularity: a measure describing how close the region is to a circular shape; Equivalent Diameter: the diameter of a circle with the same area as the measured region; Color analysis results: mean RGB, Hex, Lab color values, and color distribution results.

Users can choose to save only morphological measurements, or save both morphological and color-related measurements, depending on the purpose of the analysis. Before exporting the results, users are advised to check the Image Path, Label Name, and Shape Type fields in the Results table to ensure that each record corresponds to the correct image and annotation region. This helps avoid sample-matching errors in downstream statistical analysis.

### 7. Label Management

In BioIMA, a label is the basic unit used to store a target region or measurement object in an image. Polygon, rectangle, line, and angle annotations created manually by the user, as well as masks generated and accepted through SAM-based segmentation, can all be displayed as labels in the Labels panel. The Labels panel is used to manage all annotation objects in the current image. Users can select, rename, change the color of, or delete labels in this panel, and can also use the Mask Preview panel to check whether the currently selected label corresponds to the correct target region. Good label management helps ensure the accuracy and traceability of subsequent measurement results. This is especially important when multiple target regions are present in the same image. In such cases, users are advised to give different labels clear names, so that each measurement record can be correctly identified in the Results table and the exported CSV file.

#### 7.1 Creating a Label

Users can create labels in two ways: manual annotation and SAM-based automatic annotation.

When creating a label manually, users can select tools such as Polygon, Rectangle, Line, or Angle, and draw the target region or measurement object in the image display area. After the drawing is completed, the software saves the object as a new label and displays it in the Labels panel on the right. When creating a label through SAM, users first need to generate a mask using point prompts or box prompts. After confirming the segmentation result, users can click Accept Mask, and the software will convert the current mask into a label. The accepted mask label can be selected, previewed, edited, measured, and exported in the same way as a manually created annotation. Common label types include: Polygon: a polygonal region suitable for irregular target areas; Rectangle: a rectangular region suitable for regular areas or local image selection; Line: a line object used for length measurement; Angle: an angle object used for angle measurement; SAM mask label: a label generated from a SAM segmentation result, suitable for target regions with complex boundaries or regions that would be time-consuming to trace manually. After creating a label, users are advised to check its boundary position and name to ensure that the label corresponds to the intended target region.

##### 7.2 Selecting a Label

All labels created in the current image are displayed in the Labels panel on the right. Users can select a label by clicking an item in the Labels list. After a label is selected, the software displays the position and boundary of that label in the image display area. At the same time, the Mask Preview panel shows a local preview of the currently selected label, helping users confirm whether the target region is correct. Selecting a label is usually used for the following operations: viewing the position of the current label; checking whether the label covers the correct target region; renaming the label or changing its color; editing the label boundary or vertices; measuring the current label; deleting labels that are no longer needed.

Before clicking Measure, users should first confirm that the currently selected label is correct. If the wrong label is selected, the measurement result may correspond to an incorrect region and affect downstream data analysis.

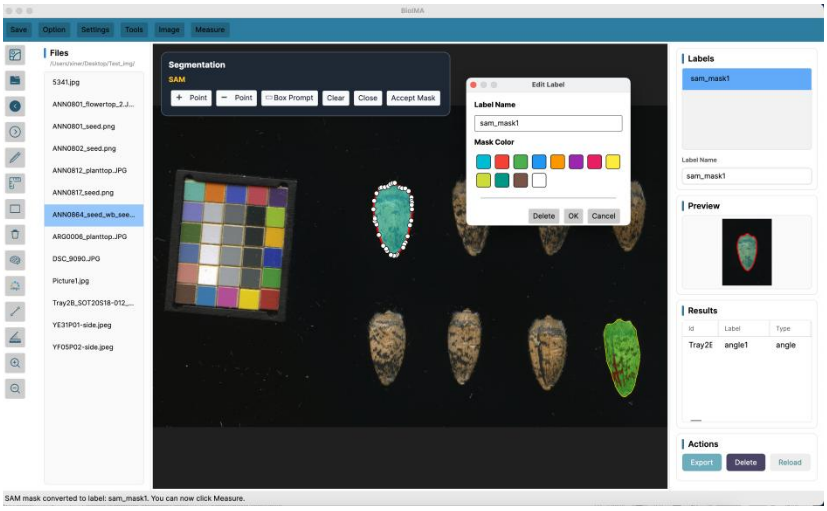

##### 7.3 Renaming a Label

To make label management and later export easier, users can rename labels. Clear label names help users identify different samples, regions, or measurement objects in the Results table. To rename a label, select the label to be renamed in the Labels panel; enter a new name in the label name editing box; after confirming the change, the Labels list will update the label name; this label name will also be recorded in subsequent measurement results. If multiple target regions are present in the same image, it is recommended to use a consistent naming rule. For example, labels can be named according to sample ID, organ name, or region number. This makes data organization and statistical analysis easier after exporting the CSV file.

##### 7.4 Changing Label Color

BioIMA supports assigning different colors to labels, allowing users to distinguish multiple annotated regions in the image. Changing the label color does not modify the original image or affect the measurement results; it only changes the display appearance in the interface. Users can select a label in the Labels panel and then change its display color using the color setting function. After the color is changed, the label boundary or overlay in the image display area will be shown in the new color. Changing label color is useful when multiple target regions are present in the same image; when different types of annotations need to be distinguished; when the boundary needs to be more visible on the image; or when a specific label needs to be highlighted in screenshots or result presentations. Color settings are mainly used for visual distinction and do not affect area, length, angle, or color analysis results.

##### 7.5 Deleting a Label

If a label was drawn incorrectly, if the segmentation result is not satisfactory, or if the label is no longer needed, users can delete it. To delete a label, select the label to be removed in the Labels panel; click the Delete button; after confirmation, the label will be removed from both the Labels list and the image display area.

Before deleting a label, users are advised to confirm that the currently selected object is correct, to avoid accidentally deleting a completed annotation. For results that have already been measured and saved in the Results panel, deleting the label may not automatically remove the saved measurement record. Therefore, if users delete an incorrect label, they should also check whether the corresponding incorrect measurement record exists in the Results panel and delete it or re-export the results if necessary.

If a SAM segmentation result is still only a temporary preview and Accept Mask has not been clicked, users can usually use Clear to remove the current prompt points or temporary mask. If the mask has already been accepted as a label, users need to select the corresponding label in the Labels panel and then delete it.

##### 7.6 Checking a Label in the Mask Preview Panel

The Mask Preview panel is used to display a local preview of the currently selected label. After a user selects a label in the Labels panel, the Mask Preview panel shows the corresponding image region, boundary, or mask overlay for that label.

The Mask Preview panel helps users quickly check whether the currently selected label corresponds to the correct target; whether the label boundary fully covers the target region; whether the SAM mask contains missing or extra segmented regions; whether the manually edited boundary is reasonable; and whether the label needs further modification before measurement.

For manually drawn polygon or rectangle labels, the Mask Preview panel mainly displays the local image region and boundary of the label. For SAM-generated mask labels, the Mask Preview panel can display the mask-covered region, helping users determine whether the segmentation result is accurate. BioIMA currently supports accepting a SAM mask as a label and checking the region through preview before measurement.

If the Mask Preview panel is empty, it may be because no label is currently selected, or because the selected object does not contain a region that can be previewed. Users can first select an existing label in the Labels panel and then view the corresponding preview.

Before formal measurement, users are encouraged to check the Mask Preview panel. This can reduce measurement errors caused by selecting the wrong label, incomplete boundaries, or inaccurate masks.

### 8. SAM-Based Segmentation

The SAM-based segmentation function is used to help users quickly and automatically obtain a mask of the target region. Compared with manually drawing a polygon, SAM reduces the need to trace the target boundary point by point, making it especially useful for biological images with complex boundaries, irregular shapes, or targets that are time-consuming to annotate manually. In BioIMA, SAM segmentation uses an interactive prompting approach. Users can indicate the target region to be segmented by using positive point prompts, negative point prompts, or box prompts. The software generates a mask based on the prompt information and displays it on the image as a semi-transparent overlay. After confirming the segmentation result, users can click Accept Mask to save the current mask as a label. The saved label can then be previewed, edited, measured, and exported. The general workflow of SAM-based segmentation is as follows: load the SAM model → open an image → wait for image encoding to complete → add positive points, negative points, or a box prompt → view the mask overlay → adjust the prompts and re-segment if needed → click Accept Mask to generate a label → check, edit, and measure the label.

#### 8.1 Loading the Model

Before using SAM-based segmentation, the corresponding segmentation model needs to be loaded by the software. The current version includes a built-in model, so users can usually enter the SAM segmentation function directly without downloading or selecting an additional model file. Depending on the software version, if users wish to use a custom model, they can select the corresponding model file through the model loading function. To load the model, start BioIMA, open the image to be analyzed, and click SAM, or the related model selection button. If the software includes a built-in model, wait for the model to load automatically. If a model needs to be loaded manually, select the corresponding SAM model file. After the model has been loaded, users can start image segmentation. Model loading may take some time, depending on the model size, computer performance, and the size of the current image. Once loading is complete, the software will enter the segmentation-ready state.

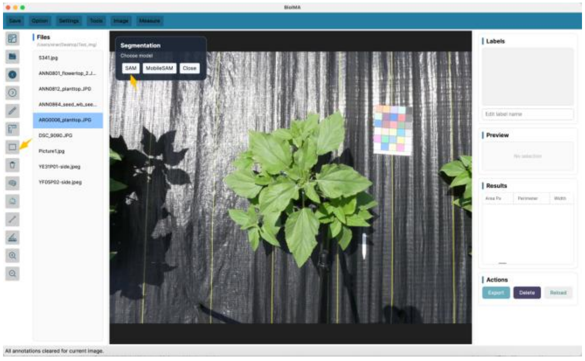

##### 8.2 Image Encoding and Model Preparation

SAM-based segmentation usually includes two main steps: image encoding and prompt decoding. Image encoding means that the software first extracts features from the current image to prepare for later interactive segmentation. Prompt decoding means that the software generates a mask for the target region based on the points or box provided by the user. After the user opens an image and starts the SAM function, the software will encode the current image. Before image encoding is completed, the segmentation function may be temporarily unavailable. Users should wait until the model is ready before adding prompt points or a prompt box. In general, users only need to pay attention to the interface status: Model loading means waiting for the model to be ready; Image encoding means waiting for feature extraction of the current image to complete; Ready / Embedding Ready means that users can start adding prompts and generating a mask.

If the user switches to another image, the software needs to encode the new image again. For high-resolution images, encoding may take longer. Users are advised to wait until the current image is ready before performing segmentation.

##### 8.3 Positive Point Prompt Segmentation

A positive point prompt is used to tell the software that “this area belongs to the target region.” When the user clicks inside the target object, SAM generates a possible target mask based on the image information around that point.

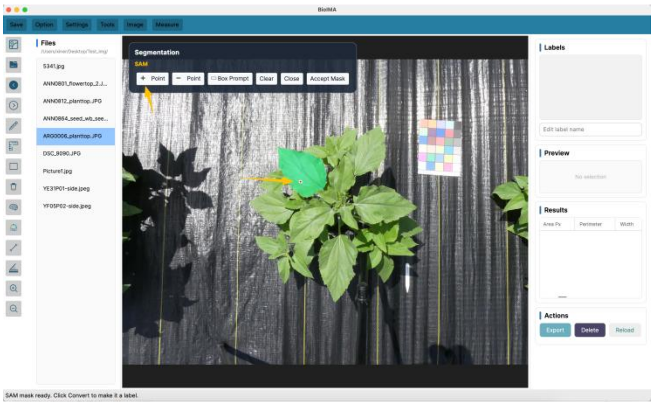

To use positive point prompt segmentation, select the SAM segmentation function, choose the + Point or positive point prompt tool, and click a point inside the target region. The software will generate and display the corresponding mask overlay. Users should then check whether the mask covers the target region. If needed, users can continue adding positive point prompts or combine them with negative point prompts to refine the result. Positive point prompts are usually suitable for images where the target region is clear and visually distinct from the background. If the mask is incomplete after the first click, users can add more positive points inside the missing parts of the target region to help the software expand the segmentation area.

When using positive point prompts, users should click a representative position inside the target region and avoid clicking on edges, shadows, or background areas.

##### 8.4 Negative Point Prompt Segmentation

A negative point prompt is used to tell the software that “this area does not belong to the target region.” When the mask generated by SAM includes extra background, adjacent objects, or unwanted regions, users can add negative point prompts to these incorrect areas to help the software exclude them. The operation steps are as follows: 1. Based on the existing mask, select the - Point or negative point prompt tool; 2. Click a point in the area that should not be included; 3. The software regenerates the mask based on the negative point prompt; 4. Check whether the extra region has been removed; 5. If incorrect segmentation remains, continue adding negative point prompts.

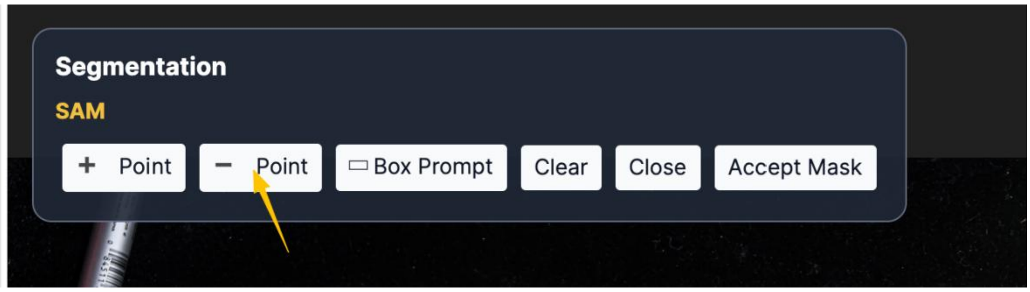

Negative point prompts are commonly used in the following situations: the mask includes background; the mask is connected to adjacent samples; the mask covers structures that do not need to be analyzed; extra regions appear near the target boundary.

Positive and negative points can be used together. In general, users can first use positive points to specify the target region and then use negative points to exclude incorrect areas until the mask coverage meets the analysis needs.

##### 8.5 Box Prompt Segmentation

A box prompt is used to specify the approximate range of the target with a rectangular box. Users can draw a box around the target region in the image, and the software will prioritize detecting and generating the corresponding mask within that box.

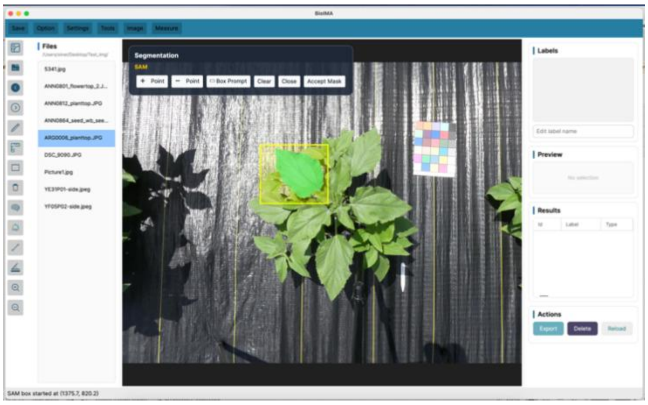

The operation steps are as follows: 1. Select the SAM segmentation function; 2. Select Box

Prompt or the box prompt tool; 3. Use the right mouse button, Drag the mouse in the image to draw a box around the target region; 4. After releasing the mouse, the software generates a mask overlay; 5. Check whether the mask matches the target boundary; 6. If needed, further refine the result using positive or negative point prompts.

Box prompts are suitable when the target region is relatively large, when the boundary is relatively clear, or when users want to quickly limit the segmentation range. For images with adjacent samples or complex backgrounds, box prompts can reduce the possibility of SAM segmenting the wrong region.

When drawing the prompt box, it is recommended that the box slightly contains the complete target region, but does not include too much unrelated background or adjacent targets. If the box is too large, incorrect segmentation may increase; if the box is too small, the target boundary may be incomplete.

##### 8.6 Clearing Prompts and Re-segmentation

If the current prompt points, prompt box, or generated mask does not meet expectations, users can use the Clear function to remove the current temporary segmentation result and add new prompts. Clear is commonly used in the following situations: the prompt point was placed incorrectly; the prompt box range is not appropriate; the generated mask differs greatly from the target region; the user needs to restart segmentation for the current target; the user wants to switch to another target region for segmentation.

The operation steps are as follows: 1. Click Clear or the clear button; 2. The current temporary prompt points, prompt box, or mask overlay will be removed; 3. Select positive point, negative point, or box prompt again; 4. Generate a new mask and check the result again.

It should be noted that Clear is usually used to remove temporary segmentation results that have not yet been accepted. If the user has already clicked Accept Mask and saved the mask as a label, the label will appear in the Labels panel. In this case, if the user wants to delete it, they should select the corresponding label in the Labels panel and click Delete. Alternatively, users can use Reload to remove all labels currently displayed on the image, including the ruler that has already been set.

##### 8.7 Accepting a Mask and Generating a Label

When users confirm that the current SAM mask covers the correct target region, they can click Accept Mask. The software will save the current mask as a new label and display it in the Labels panel.

The operation steps are as follows: 1. Generate a mask using positive point prompts, negative point prompts, or a box prompt; 2. Check the mask overlay on the image; 3. After confirming that the mask covers the target region, click Accept Mask; 4. The software converts the current mask into a label; 5. The new label will appear in the Labels panel; 6. Users can select this label, view it in the Mask Preview panel, and perform measurement.

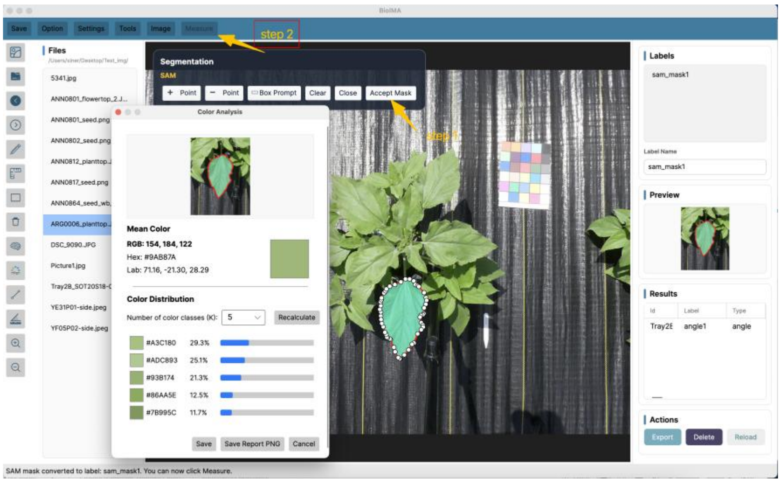

The accepted SAM mask label can be managed like a regular manually created label, including selecting, renaming, changing color, deleting, previewing, and measuring.

Users are advised to carefully check whether the mask fully covers the target region and whether it includes extra background or adjacent objects before clicking Accept Mask. If the segmentation result is not satisfactory, users should first adjust it using positive points, negative points, or a box prompt, and then accept it as a label.

##### 8.8 Editing a SAM-Generated Label

A SAM-generated label can still be checked and edited after it has been accepted. If users find that the boundary is not fully accurate, they can modify the label using vertex editing or boundary adjustment functions.

Common cases that may require editing include: the mask misses part of the target boundary; the mask includes a small amount of background; the mask is connected to adjacent objects; the target boundary is locally inaccurate; users want to manually correct the analysis region according to their research criteria. To edit the label, users can first select the corresponding label in the Labels panel, and then view its boundary and control points in the image display area. After adjustment, users can check the local preview of the label again in the Mask Preview panel.

Editing is important for SAM-generated labels. If the user does not modify the boundary, the software will try to preserve the measurement accuracy of the original SAM mask. If the user manually edits the boundary, subsequent measurements will be calculated based on the edited region. This design preserves the convenience of SAM-basedsegmentation while still allowing users to manually correct the region according to the actual image.

##### 8.9 Measurement Method for SAM Masks

After a SAM mask is accepted as a label, it can be measured in the same way as a manual annotation. Users only need to select the corresponding SAM mask label in the Labels panel and click Measure. The software will then calculate the morphological and color-related measurements for that region.

The measurement results of a SAM mask label may include: Area: real-world area, which depends on scale calibration; Area Px: pixel area; Perimeter: perimeter of the region; Width / Height: width and height of the target region; Aspect Ratio: width-to-height ratio; Circularity: circularity of the region; Equivalent Diameter: equivalent diameter; Mean RGB / Hex / Lab: mean color values of the target region; Color Distribution: the proportions of different color classes within the target region.

For SAM mask labels that have not been edited, the software uses the original mask for measurement by default, in order to preserve the accuracy of the segmentation boundary as much as possible. For SAM labels whose boundaries have been manually edited, the software recalculates the measurement results based on the edited region. The current design of BioIMA is that a simplified boundary can be used for display and editing, while the original SAM binary mask is retained as much as possible for precise measurement; if the user edits the boundary, the edited region will be used for measurement. Before formally saving the measurement results, users are advised to check the current label in both the image display area and the Mask Preview panel. If the mask does not fully cover the target or contains obvious incorrect regions, it should be adjusted before clicking Measure to save the result.

### 9. Color Analysis

BioIMA provides color analysis functions for extracting color information from target regions. Users can perform color measurements on a rectangular selected region, a manually drawn label, or a mask label generated by SAM segmentation. Color analysis results can be used to describe the overall color of a sample, compare color differences among individuals, or quantify the proportions of different color classes within a target region.

Color analysis mainly includes two types of results. The first type is mean color, including RGB, Hex, and Lab color values. The second type is color proportion analysis, namely Color Distribution, which is used to calculate the major color classes within the target region and their proportions. BioIMA currently supports mean color calculation, K-means-based color distribution analysis, K selection, recalculation, and PNG export of color analysis reports.

#### 9.1 Color Correction

The Color Correction function is located under the Image menu in the top menu bar.

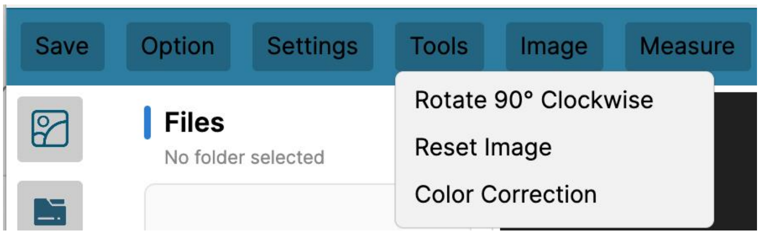

**Color Correction** is used to reduce color bias caused by lighting, camera settings, or scanning conditions. This function applies a basic white balance or gray reference correction to the current image based on a white or gray reference region selected by the user. Color correction is suitable for use before formal color measurement, especially when users need to compare color differences among different samples.

Color correction is based on a user-selected neutral reference region. This region should be an area that is expected to be close to white or gray in the image, such as a gray card, white paper, white scale bar, scale bar, or neutral color standard block. The software calculates the mean RGB value of the selected reference region and uses it to adjust the RGB channels of the whole image, thereby reducing overall color cast.。

The operation steps are as follows: 1. Click Image in the top menu bar; 2. Select Color Correction; 3. Click the first corner of a white or gray reference region in the image; 4. Click the opposite corner of the reference region to define the reference box; 5. BioIMA calculates the mean RGB value of the selected reference region; 6. The software applies color correction to the whole image based on the selected reference region; 7. Subsequent color measurements and Color Distribution analyses will be based on the corrected image.

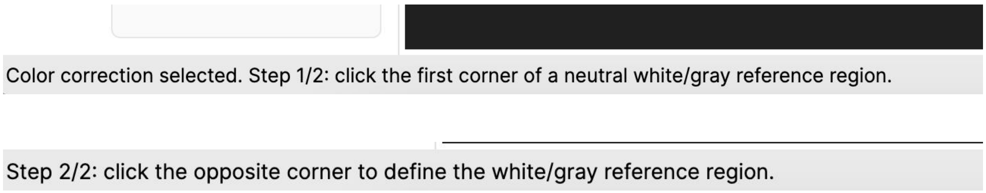

Recommended reference regions include: neutral gray card; white or gray color standard block; white paper; white scale bar; neutral light-gray background. Reference regions that should be avoided include: clearly colored regions; black background; strong shadow areas; reflective areas; overexposed white highlights; dirty or strongly textured areas.

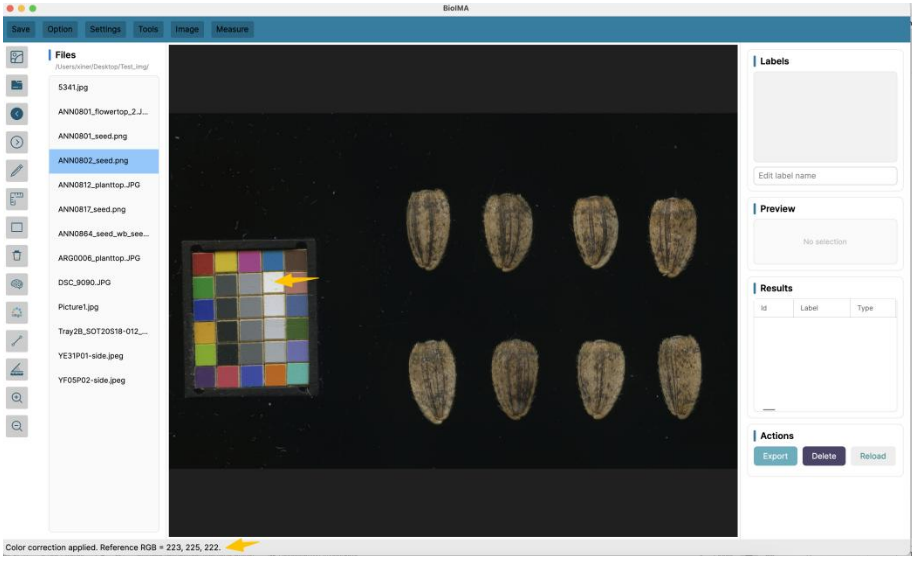

It should be noted that Color Correction changes the image pixels used for color measurement, but does not change the spatial positions of labels or masks. Therefore, existing manual labels and accepted SAM masks remain aligned with the corrected image. For SAM-based analysis, SAM is mainly used to define the measurement region, while color values are extracted from the current corrected image during measurement. Therefore, accepted SAM masks can still be used for corrected color analysis.

##### 9.2 Mean Color Measurement

Mean color measurement is used to calculate the average color of all pixels within the target region. Users can select a region using the Color Box tool, or choose an existing label and perform color analysis on that region. Mean color measurement is suitable for comparing the overall color intensity of different samples; recording the representative color of an organ or region; extracting the mean color of petals, seeds, leaves, or patterned regions; and providing standardized color parameters for downstream statistical analysis.

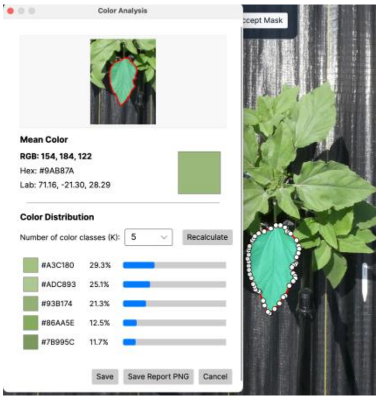

Mean color results usually include: Mean RGB: the average red, green, and blue channel values of the target region; Hex: the hexadecimal color value converted from the mean RGB value; Lab: the color value of the target region in the CIE Lab color space. It should be noted that mean color represents the overall color tendency of the selected region. If the target region contains large internal color variation, such as both dark patterns and a light background, using only the mean color may obscure local differences in color composition. In this case, the Color Distribution function can be used to further analyze color proportions.

##### 9.3 RGB, Hex, and Lab Color Values

The color analysis results in BioIMA include three commonly used color representations: RGB, Hex, and Lab. RGB represents color using three channels: red, green, and blue. Each channel is usually represented by a numerical value, and a higher value indicates a stronger contribution from that color channel. For example, the Mean RGB of a region can be shown as: Mean RGB: 120, 85, 60. Hex is a hexadecimal color code converted from RGB values and is commonly used in web design, graphic design, and color recording. For example: Hex: #78553C. Lab is a color space that is closer to human visual perception and is commonly used for color difference analysis. Lab usually contains three values: L: lightness, representing how light or dark the color is; a: the red-green axis, where positive values indicate red and negative values indicate green; b: the yellow-blue axis, where positive values indicate yellow and negative values indicate blue.

In biological image analysis, Lab color values are often used to compare color differences among samples. Compared with RGB, Lab is better suited for describing lightness, red-green variation, and yellow-blue variation, making it more interpretable for studies of plant color, seed color, petal color, and related traits.

##### 9.4 Color Box Region Color Analysis

The Color Box tool 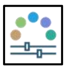 is used to quickly select a rectangular region and perform color analysis. This function is suitable for analyzing the mean color or color composition of a local region, such as a patch, a local tissue area, or a background region in the image. The operation steps are as follows: 1. Open the image to be analyzed; 2. Select the Color Box tool from the left toolbar; 3. Drag the mouse in the image display area to select the region to be analyzed; 4. After releasing the mouse, the software opens the Color Analysis window; 5. View the ROI preview, Mean RGB, Hex, Lab, and Color Distribution in the window; 6. If needed, adjust the number of color classes, K, and click Recalculate; 7. Click Save to save the result to the Results panel; 8. Click Save Report PNG to export the color analysis report.

The Color Box tool is suitable for quickly analyzing regular rectangular regions. If users want to perform color analysis along the true boundary of a target, such as analyzing only one petal or one seed, it is recommended to first generate a label using Polygon, Rectangle, or SAM, and then perform color analysis on that label.

##### 9.5 Label Region Color Analysis

In addition to Color Box, BioIMA can also perform color analysis on existing labels. A label can be created from manual annotation or from a mask accepted after SAM-basedsegmentation. Compared with Color Box, label region color analysis is more suitable for samples with irregular target boundaries.

The operation steps are as follows: 1. Create a target region label using Polygon, Rectangle, or SAM; 2. Select the label to be analyzed in the Labels panel on the right; 3. Check the Mask Preview to confirm that the label covers the target region; 4. Click Measure or the related color analysis button; 5. View the mean color and color proportion results in the Color Analysis window; 6. Adjust the K value and recalculate if needed; 7. Click Save to save the color analysis results.

For a manually drawn polygon label, the software calculates color values based on the area inside the polygon. For a SAM mask label, the software calculates color values based on the region covered by the mask. Therefore, SAM labels are usually more suitable for analyzing regions with complex or irregular boundaries. Before saving the results, users are advised to check the Mask Preview and the ROI / mask preview in the Color Analysis window to make sure that the analyzed region does not include obvious background or unrelated objects.

##### 9.6 Color Distribution Analysis

Color Distribution is used to analyze the major colors within a target region and the proportion of each color. This function is based on K-means clustering, which divides the pixels in the target region into several color classes and calculates the percentage of each class.

For example, a seed region may be divided into five color classes: Cluster 1: 42.3%; Cluster 2: 27.8%; Cluster 3: 15.4%; Cluster 4: 9.1%; Cluster 5: 5.4%. These results represent the relative proportions of different color classes within the target region. Color Distribution is suitable for analyzing the color composition of seed or fruit surfaces; comparing color proportions in petals, leaves, or patterned regions; describing the distribution of dark, light, or intermediate colors within a target region; and extracting color composition features for statistical analysis.

It should be noted that Color Distribution is a color proportion analysis. It divides a region into different color classes based on pixel color similarity. Users need to interpret the biological meaning of each color class according to the sample characteristics and research purpose.

##### 9.7 Selecting the Number of Color Classes, K

K represents the number of color classes used in color clustering. For example, K = 5 means that the software will divide the pixels in the target region into five major color classes.

In BioIMA, users can select the K value in the Color Analysis window. By default, the software uses a preset K value of 5 for analysis, but users can adjust K according to the image characteristics and research needs. For example, if the target region mainly contains a light background and dark patterns, users can try K = 2 or K = 3. If the target region contains multiple color levels, such as red, yellow, brown, and shadow areas, users can try K = 4 – 6. It is recommended to use the same K value for samples in the same batch to ensure that the color proportion results are comparable across samples.

##### 9.8 Recalculating Color Proportions

After changing the K value, users need to click Recalculate to recompute the color proportions. The software will perform color clustering again based on the new K value and update the Color Distribution results. The operation steps are as follows: 1. View the current color analysis results in the Color Analysis window; 2. Select a new number of color classes, K; 3. Click Recalculate; 4. The software recalculates the color classes and their corresponding proportions; 5. Check the updated color distribution results; 6. After confirmation, click Save to save the results.

Recalculation does not modify the original image and does not automatically overwrite exported files. The current color analysis results are saved to the Results panel only after the user clicks Save. When adjusting the K value, users can observe whether the color classes generated under different K values are reasonably interpretable. For example, some K values may split the same color region into too many classes, while others may merge different colors into the same class. Users should choose an appropriate K value based on the image content and research purpose.

##### 9.9 Saving Color Analysis Results

After completing color analysis, users can save the results to the Results panel or export a color analysis report as a PNG file. After clicking Save, the current color analysis result will be saved as a record in the Results panel. The saved content usually includes Image Path: the path of the current image; Label Name: the name of the current analysis region or label; Shape Type: the type of analysis region, such as box, polygon, or mask; Mean Rgb: the mean RGB value; Hex Color: the Hex code corresponding to the mean color; Lab: the mean Lab color value; Color ClusterK: the number of color classes, K; Dominant Color Hex: the Hex code of the dominant color class; Color Distribution: the proportions of each color class.

After clicking Save Report PNG, the software exports the current Color Analysis window as a PNG report. The report usually contains the ROI / mask preview, mean color information, and color distribution results. It is suitable for result recording, presentation, or later checking. Before saving color results, users are advised to confirm whether the analysis region is correct; whether the Mask Preview or ROI preview contains the target region; whether extra background is included; whether the K value is suitable for the current sample; and whether Recalculate has been clicked to update the results. After saving, users can check the record in the Results panel and export the CSV file after completing a batch of image analyses.

### 10. Viewing and Exporting Results

The measurement results in BioIMA are saved in the Results table. After users complete region measurement, line measurement, angle measurement, or color analysis, the result can be saved as a record and viewed in the Results panel. After completing a batch of image analyses, users can export the Results table as a CSV file for downstream statistical analysis, plotting, or integration with other data.

In addition to CSV files, BioIMA also supports exporting color analysis reports as PNG files. A color analysis report can record the preview of the current analysis region, mean color information, and color distribution results, making it useful for later checking and presentation. The current version of BioIMA supports the Results DataGrid, CSV export, saving color analysis results, and exporting color report PNG files.

##### 10.1 Results Table

The Results table is used to display the measurement records saved by the user. Each time a user completes a measurement and clicks save, the software adds a new row to the Results panel. This record usually includes information such as image path, label name, region type, morphological measurements, and color analysis values.

The Results panel helps users view the measurement records that have been completed; check the image and label corresponding to each record; confirm whether area, length, angle, and color results have been saved successfully; check whether the data are complete before exporting the CSV file; and organize the result table for downstream statistical analysis.

During batch analysis, users are advised to check whether the correct record has been added to the Results table after measuring each sample or label. This is especially important when multiple labels are present in the same image. Users should confirm that the Image Path, Label Name, and Shape Type of each record are correct.

##### 10.2 Description of Morphological Measurement Fields

The Results table contains several morphological measurement fields. Different types of labels generate different measurement results. For example, polygon, rectangle, and SAM mask labels usually generate area, perimeter, and shape-related measurements; line labels generate length results; and angle labels generate angle results. The morphological measurement fields are as follows: Id: the record number of the result; Image Path: the image path corresponding to the current measurement result; Label Name: the label name of the current measurement object; Shape Type: the type of measurement object, such as polygon, rectangle, line, angle, or mask; Area: the area in real-world units. This result requires scale calibration; AreaPx: the pixel area, namely the number of pixels contained in the target region; Perimeter: the perimeter of the target region; Width: the width of the target region; Height: the height of the target region; Aspect Ratio: the width-to-height ratio, usually used to describe the shape proportion of the target region; Circularity: circularity, used to describe how close the target region is to a circular shape; Equivalent Diameter: the equivalent diameter, namely the diameter of a circle with the same area as the target region; Line Length: the line segment length in real-world units. This result requires scale calibration; LinePx: the pixel length of the line segment; Angle Degree: the angle value, in degrees.

If no scale has been set, fields related to real-world units may be empty or shown as NA, but pixel area, pixel length, and other pixel-based results can still be saved normally. If users need to compare real-world area or real-world length across different images, they should complete scale calibration before measurement.

##### 10.3 Description of Color Measurement Fields

When users perform color analysis, color-related fields are saved in the Results table. These fields can be used to describe the mean color and color composition of the target region.

The color measurement fields are as follows: Mean Rgb: the mean RGB color value of the target region; Hex Color: the hexadecimal color code converted from the mean RGB value; Lab: the mean Lab color value of the target region; MeanR: the mean red channel value; MeanG: the mean green channel value; MeanB: the mean blue channel value; LabL: the lightness value in the Lab color space; LabA: the red-green axis value in the Lab color space; LabB: the yellow-blue axis value in the Lab color space; Color ClusterK: the number of color classes, K, used in Color Distribution analysis; Dominant ColorHex: the Hex color value corresponding to the color class with the highest proportion in the target region; ColorDistribution: the color classes and their proportions.

Among these fields, Mean Rgb, Hex Color, and Lab represent the overall mean color of the target region. Color ClusterK, Dominant ColorHex, and Color Distribution are used to record the color proportion analysis results. For example, Color Distribution may be shown as: Cluster 1: 42.3%; Cluster 2: 27.8%; Cluster 3: 15.4%; Cluster 4: 9.1%; Cluster 5: 5.4%.

This result indicates that the target region has been divided into five major color classes, and the proportion of each color class has been calculated. Users can interpret the biological meaning of different color classes according to their research purpose.

##### 10.4 Exporting a CSV File

After completing measurement, users can export the Results table as a CSV file. The CSV file can be used for downstream analysis in Excel, R, Python, or other statistical software. After completing measurement, users can export the Results table as a CSV file. The CSV file can be used for downstream analysis in Excel, R, Python, or other statistical software. The operation steps are as follows: 1. Confirm that the Results panel already contains the measurement records to be exported; 2. Click Export CSV or the export button; 3. Select a save location in the pop-up save window; 4. Enter a file name; 5. Click Save, and the software will export the Results table as a CSV file. The exported CSV file usually contains all fields in the Results table, such as image path, label name, morphological measurement results, color values, and color proportion results. Users can select specific columns for downstream analysis as needed.

The CSV file is the main data file for downstream statistical analysis. Users are advised to export it promptly after completing a batch of image analyses and keep the original CSV file to avoid losing traceability after later modifications.

##### 10.5 Exporting a Color Analysis Report as PNG

In addition to exporting the CSV results table, BioIMA also supports exporting the Color Analysis window as a PNG report. The color analysis report usually contains the preview of the current analysis region, mean color, RGB / Hex / Lab values, and Color Distribution results.

The operation steps are as follows: 1. Perform color analysis using Color Box or a label region; 2. Check the ROI / mask preview in the Color Analysis window; 3. View the Mean RGB, Hex, Lab, and Color Distribution results; 4. If the number of color classes, K, needs to be adjusted, select a new K value and click Recalculate; 5. After confirming the results, click Save Report PNG; 6. Select a save location and enter a file name; 7. Click Save to export the PNG report. The color analysis report PNG is suitable for recording the color analysis process of a single sample or region; saving the ROI or mask preview; presenting mean color and color proportion results; serving as supplementary information in papers, reports, or experimental records; and later checking whether the color analysis region was correct.

It should be noted that the PNG report is mainly used for visual documentation, while the CSV file is the main data file for statistical analysis. Users are advised to save both the CSV results and the necessary PNG reports as needed, so that data analysis and result traceability can both be supported.

##### 10.6 Recommendations for Saving Result Files

To ensure that analysis results are clear and traceable, users are advised to create a separate result folder for each project and save different types of output files separately.

The recommended file structure is as follows:

```
Project_Name/
 ├── images/
 ├── results/
 └── color_reports/
```

In this structure, images stores the original image files; results stores the exported CSV measurement results; color_reports stores the exported color analysis report PNG files.

Users are advised not to directly modify the original image files. All measurement results should be saved through the Results panel and exported as CSV files. Color analysis reports can be saved as supplementary records in the color_reports folder.

File names should also be as clear as possible, for example: results_seed_batch01.csv; results_leaf_area_2026.csv; color_report_sample_001.png; color_report_petal_left.png.

For batch image analysis, it is recommended to include the project name, sample batch, date, or analysis type in the file name. This can reduce confusion between different result versions and make later checking and statistical analysis easier.

Before formally using the results, users are advised to open the exported CSV file for a simple check and confirm that the column names, sample names, and measurement values are correct. If errors are found, users can return to BioIMA to recheck the corresponding label, scale setting, or color analysis settings, and then export the results again.

#### 11. Usage Recommendations and Notes

To obtain stable, accurate, and reproducible measurement results, users should pay attention to factors such as image quality, background conditions, scale settings, inspection of SAM segmentation results, and color analysis parameter selection when using BioIMA for image analysis. This section summarizes common usage recommendations to help users reduce measurement errors and improve result comparability.

#### 11.1 Image Quality Requirements

Image quality directly affects annotation, segmentation, and measurement results. Users are advised to keep imaging or scanning conditions as clear, stable, and consistent as possible when acquiring images.

Users should pay attention to the following points: 1. Images should be as clear as possible and avoid obvious blur or defocus; 2. The target region should appear completely in the image and should not be cropped; 3. Lighting should be as even as possible, avoiding strong reflections, shadows, or local overexposure; 4. Samples in the same batch should be captured or scanned using the same parameters whenever possible; 5. If color analysis is needed, the background, light source, and exposure conditions should be kept consistent; 6. If real-world area or real-world length is needed, the image should include a scale bar, or the image resolution should be known. For color analysis, image acquisition conditions are especially important. Different lighting, exposure, white balance, or scanning settings may affect RGB, Lab, and Color Distribution results. Therefore, if users need to compare color differences among samples, it is recommended to acquire images under consistent conditions, perform color correction, and use the same image-processing workflow as much as possible.

#### 11.2 Recommendations for Images with Complex Backgrounds

When the image background is complex, when the target and background have similar colors, or when multiple samples are touching each other, both manual annotation and SAM-assisted segmentation may be affected. In this case, users should choose appropriate tools and operations according to the image characteristics. For images with complex backgrounds, the following practices are recommended: 1. Prioritize images with clear target boundaries for analysis; 2. If the target region is regular, use Polygon or Rectangle for manual annotation; 3. If the target boundary is complex, use SAM-assisted segmentation to reduce manual tracing; 4. When using SAM, first use Box Prompt to limit the target range; 5. If the mask includes background or adjacent targets, use negative point prompts to exclude them; 6. Before clicking Accept Mask, carefully check the mask overlay; 7. After saving the label, confirm the region again through the Mask Preview panel.

If the target region is very similar to the background, SAM segmentation results may be unstable. In this case, users can try adding more prompt points, reducing the box prompt range, or switching to manual polygon annotation. For measurements that require high-precision boundaries, users are advised to manually inspect the segmentation result and make necessary edits after segmentation.

##### 11.3 Notes on Scale Setting

Scale calibration affects area and length results in real-world units. If the scale is set incorrectly, all subsequent measurements in real-world units will be affected. Therefore, before measuring real-world area or real-world length, users should carefully check the scale setting.

When setting the scale, users should note the following: 1. The ruler line should overlap as closely as possible with the scale bar or known-length region in the image; 2. The starting point and endpoint should be accurately placed at the two ends of the known length; 3. When entering the real-world length, users should confirm that both the value and unit are correct; 4. Do not confuse units such as mm, cm, and μm; 5. If different images have different imaging distances, scanning resolutions, or zoom levels, the scale should be set separately for each image; 6. If a batch of images was acquired using exactly the same parameters, the same scale setting can be used. If users only need pixel-level comparison, scale calibration is not required. However, if users need to compare real-world area or real-world length across different images or samples, scale calibration should be completed first. When measurement results appear obviously unreasonable, users should first check whether the scale was set correctly, such as whether the ruler line was drawn in the wrong position, whether the real-world length was entered incorrectly, or whether the wrong unit was selected.

##### 11.4 Checking and Editing SAM Segmentation Results

SAM-assisted segmentation can reduce manual tracing work, but it is still an interactive segmentation method based on user prompts. Users should not use segmentation results for measurement without checking them first. After using SAM, users are advised to check the following points: 1. Whether the mask fully covers the target region; 2. Whether the mask includes extra background; 3. Whether the mask is connected to adjacent samples; 4. Whether any obvious parts of the target boundary are missing; 5. Whether there are uncovered holes inside the target region; 6. Whether the current label corresponds to the correct image and sample.

If the segmentation result is not satisfactory, users can first use Clear to remove the current temporary mask, and then add positive points, negative points, or a box prompt again. In general, positive points are used to specify the target region, negative points are used to exclude unwanted regions, and box prompts are used to limit the approximate range. After the mask is accepted as a label, users can still check and edit the label. For SAM mask labels that have not been edited, the software uses the original mask for measurement by default. If the user manually adjusts the boundary, subsequent measurements will be calculated based on the edited region. BioIMA currently supports accepting a SAM mask as a label and checking its local region through the Mask Preview panel before measurement.

##### 11.5 Recommendations for Selecting K

In Color Distribution analysis, K represents the number of color classes into which the target region is divided. The K value affects the color clustering results, so it should be selected reasonably according to the image characteristics and research purpose.

General recommendations are as follows: 1. For regions with simple colors, a smaller K value can be selected, while for regions with complex color composition, K can be increased appropriately; 2. If users only want to distinguish light and dark colors, they can try K = 2 or K = 3; 3. If the target region contains multiple color levels, users can try K = 4 – 6; 4. Samples in the same batch should use the same K value whenever possible to ensure comparability; 5. It is not recommended to blindly increase K just to obtain more complex results.

If K is too small, color differences may be oversimplified. If K is too large, the same color region may be divided into multiple classes, making the results more complex and harder to interpret. Before formal analysis, users can test different K values on several representative images and check whether the color classes match expectations. After determining an appropriate K value, the same setting should be used for the entire batch of samples.

##### 11.6 Notes on Color Distribution

Color Distribution is an analysis of color composition proportions. Its core function is to divide pixels within the target region into several classes according to color similarity and calculate the proportion of each color class. For example, in a petal region, Color Distribution can tell users how much of the region is light-colored, how much is dark-colored, how much each color class accounts for, and which color class is dominant. If the research goal is to analyze spots, users can first use Color Distribution to observe the proportions of different color classes, and then determine which classes may correspond to spotted regions based on the color class results. For more precise spot analysis, specialized spot detection or connected-component analysis methods can be further combined later. Therefore, when using Color Distribution results, users are advised to note the following: 1. It reflects color proportions and is not an automatic biological classification; 2. The same color class may represent different structures in different images; 3. Result interpretation should be combined with the original image and ROI preview; 4. Samples in the same batch should use the same K value and analysis workflow whenever possible; 5. If it is used for spot proportion analysis, the criteria for identifying color classes should be clearly described. This can help avoid over-interpreting color clustering results as specific biological structures and improve the reliability of color analysis results in downstream statistical analysis.

Users should also note that the results may be affected by image acquisition conditions, lighting, exposure, white balance, background, and color correction status. If users need to compare color proportions among samples in the same batch, it is recommended to use consistent image acquisition conditions and decide before analysis whether Color Correction should be applied uniformly. For samples in the same batch, users should keep the same color correction workflow, the same ROI or label definition method, and the same K value whenever possible.

#### 12. Frequently Asked Questions

This section summarizes common issues that users may encounter when using BioIMA and provides suggested solutions. If users experience problems such as images failing to open, model loading failure, unsatisfactory SAM segmentation, missing real-world units in measurement results, abnormal CSV display, or difficulty interpreting color results, they can first refer to this section for troubleshooting.

##### 12.1 Image Cannot Be Opened

If an image cannot be opened properly, the issue may be related to the image format, file path, file corruption, or software version. Users can check the following items in order: 1. Confirm that the image file exists and has not been moved or deleted; 2. Confirm that the image format is a common format, such as PNG, JPG, JPEG, BMP, or TIFF; 3. Try copying the image to a folder with an English path or a shorter path before opening it again; 4. Check whether the file name contains special symbols; 5. Confirm that the image file is not corrupted by opening it first with the system image viewer; 6. If the image size is too large, try using a device with more memory, or first reduce the size of a copy of the image for testing.

If a single image can be opened, but no files are displayed after opening a folder, it may be because the folder does not contain recognizable image formats, or because the images are stored in deeper subfolders. It is recommended to place the images to be analyzed directly in the same folder and use clear and consistent file names.

##### 12.2 Model Loading Failure

If model loading fails, the problem may be caused by missing model files, an incorrect file path, an incompatible model format, or incompatibility between the software version and the model file. Users can check the following items: 1. The current software version includes a built-in model, so users should confirm that the files in the installation directory have not been moved or deleted; 2. If using a custom model, confirm that the correct model file has been selected; 3. Confirm that the model file format is supported by the current software version; 4. Avoid placing model files in folders with special characters or overly long paths; 5. Restart the software and load the model again; 6. If the model file is large, loading may take some time.

If the software indicates that the model is not ready yet, users should wait until both model loading and image encoding are completed before adding point prompts or box prompts. For high-resolution images, image encoding may take longer. If the problem continues, users are advised to record the error message, software version, system information, and model file name, and then contact the author.

##### 12.3 Incomplete SAM Segmentation Results

Incomplete SAM segmentation results are usually related to the prompt point position, prompt box range, target boundary clarity, background complexity, or image quality. SAM is an interactive assisted segmentation tool, and the segmentation result should be checked and corrected by the user when necessary. If the mask does not fully cover the target region, users can try the following: 1. Add more positive point prompts inside the missing parts of the target region; 2. Use Box Prompt to select the complete target region; 3. Make sure that the prompt box contains the full target, but does not include too much background; 4. Zoom in on the image and select the prompt points again.

If the mask includes extra background or adjacent targets, users can try the following: 1. Add negative point prompts in the extra regions; 2. Reduce the range of the Box Prompt; 3. Click Clear to remove the current result and segment again; 4. Carefully check the mask overlay before clicking Accept Mask; 5. After accepting the mask as a label, check the region again through the Mask Preview panel. After a SAM mask is accepted as a label, users can still edit and measure it. BioIMA currently supports SAM point / box prompts, converting an accepted mask into a label, and measuring SAM mask labels.

##### 12.4 Measurement Results Do Not Have Real-World Units

If the Results table only contains pixel area or pixel length, but does not contain area or length in real-world units, this is usually because the scale has not been set, or the scale setting is incomplete. Users can check the following items: 1. Whether the Ruler tool has been used to draw a ruler line; 2. Whether the real-world length corresponding to the ruler line has been entered; 3. Whether the unit has been entered correctly, such as mm, cm, or μm; 4. Whether the ruler line corresponds to the scale bar or known-length region in the image; 5. Whether the current image has the same resolution and zoom ratio as the image used when setting the scale.

If the scale has not been set, BioIMA can still output pixel-based results, such as AreaPx and

LinePx. However, if real-world area or real-world length is needed, scale calibration must be completed first. If the real-world unit results are obviously too large or too small, users should first check whether the ruler line length, real-world length input value, and unit are correct.

##### 12.5 CSV Columns Are Misaligned

If the exported CSV file shows misaligned columns after being opened in Excel or other software, this is usually because some fields contain commas, such as the Mean RGB value: MeanRgb = 91, 78, 67. If CSV fields are not properly escaped, content containing commas may be incorrectly treated as multiple separate columns. BioIMA' s CSV export should escape fields containing commas, quotation marks, or line breaks to ensure that the exported table structure is correct.

Users can try the following methods: 1. Use Excel' s Import from Text/CSV function to open the file instead of double-clicking it directly; 2. Confirm that the delimiter is set to comma; 3. If the columns are still misaligned, check whether fields such as MeanRgb, Lab, or ColorDistribution contain commas; 4. Open the CSV file with a text editor and check whether fields containing commas are enclosed in double quotation marks; 5. Use the built-in Export CSV function of the software whenever possible, and avoid manually copying content from the Results table. Under normal conditions, fields containing commas should be saved in a format similar to "91, 78, 67". In this way, when the file is read by Excel, R, or Python, the entire content will be recognized as one column.

##### 12.6 How to Interpret Color Analysis Results

Color analysis results mainly include two parts: mean color and Color Distribution. Mean color is used to describe the overall color of the target region, while Color Distribution is used to describe the proportions of different color classes within the region. Mean color fields can be understood as follows: MeanRgb: the mean red, green, and blue channel values of the target region; HexColor: the hexadecimal color code corresponding to the mean color; Lab: the mean color value of the target region in the Lab color space. Color Distribution can be understood as the proportions of different color classes in the target region. For example: Cluster 1: 42.3%; Cluster 2:

27.8%; Cluster 3: 15.4%; Cluster 4: 9.1%; Cluster 5: 5.4%. This means that the software divides the pixels in the target region into five color classes and calculates the proportion of each class. The class with the highest proportion usually corresponds to DominantColorHex, which is the Hex code of the dominant color class. It should be noted that Color Distribution is an analysis of color composition proportions. It does not automatically determine whether a certain color class necessarily represents spots, lesions, or a specific biological structure. Users need to interpret these color classes together with the original image, ROI preview, mask preview, and research purpose.

If users want to compare a batch of samples, the following practices are recommended: 1. Use the same image acquisition conditions; 2. Use the same method for defining analysis regions; 3. Use the same K value for samples in the same batch; 4. Save CSV results for statistical analysis; 5. Save color analysis report PNG files when necessary for later checking of the analysis regions. If a color result appears unreasonable, users should return to the Color Analysis window and check whether the analysis region contains background, shadows, reflections, or unrelated structures, and then reselect the region or adjust the K value if needed.
